# Universal closed-tube barcoding for monitoring the shark and ray trade in megadiverse conservation hotspots

**DOI:** 10.1101/2022.11.30.518468

**Authors:** Andhika P. Prasetyo, Marine Cusa, Joanna M. Murray, Firdaus Agung, Efin Muttaqin, Stefano Mariani, Allan D. McDevitt

**Affiliations:** School of Science, Engineering and Environment, University of Salford, Salford, UK; Centre Fisheries Research, Ministry for Marine Affairs and Fisheries, Indonesia; Research Centre for Conservation of Marine and Inland Water Resources, National Research and Innovation Agency, Indonesia; Oceana Europe, Madrid, Spain; Centre for Environment, Fisheries and Aquaculture Science (CEFAS), Lowestoft, UK; Directorate for Conservation and Marine Biodiversity, Ministry for Marine Affairs and Fisheries, Indonesia; Wildlife Conservation Society Indonesia Program, Indonesia; School of Biological and Environmental Sciences, Liverpool John Moores University, Liverpool, UK; Department of Natural Sciences and the Environment, School of Science and Computing, Atlantic Technological University, Galway, Ireland

**Keywords:** elasmobranchs, DNA barcoding, deep learning, illegal trade, biodiversity monitoring, Indonesia

## Abstract

Trade restrictions for many endangered elasmobranch species exist to disincentivise their exploitation and curb their declines. However, the variety of products and the complexity of import/export routes make trade monitoring challenging. We investigate the use of a portable, universal, DNA-based tool which would greatly facilitate *in-situ* monitoring. We collected shark and ray samples across the Island of Java, Indonesia, and selected 28 species (including 22 CITES-listed species) commonly encountered in landing sites and export hubs to test a recently developed real-time PCR single-assay originally developed for screening bony fish. We employed a deep learning algorithm to recognize species based on DNA melt-curve signatures. By combining visual and machine learning assignment methods, we distinguished 25 out of 28 species, 20 of which were CITES-listed. With further refinement, this method can provide a practical tool for monitoring elasmobranch trade worldwide, without the need for a lab or the bespoke design of species-specific assays.

**Highlights:**

1. We applied a portable, universal, closed-tube DNA barcoding approach originally developed for bony fishes to distinguish between shark and ray species traded in Indonesia.
2. We built a deep machine learning model to automatically assign species from the qPCR fluorescence spectra produced by two barcodes
3. The model achieved 79.41% accuracy for classifying 28 elasmobranch species, despite the barcode regions being designed for teleost species
4. This tool can serve as a potent single-assay *in-situ* diagnostic tool to regulate trade operations and it will be significantly enhanced by further optimisation of the barcode regions to fit elasmobranch DNA sequence variation

## Introduction

Biodiversity is depleting more rapidly than at any time in human history. Within the last 50 years, animal species have declined by an average of almost 70% due to continued and increasing anthropogenic stressors (Bar-On et al., 2018; Leung et al., 2020). Shark and ray populations (hereafter referred to as ‘elasmobranchs’) have one of the highest extinction risks across the animal kingdom due to fishing pressure, whether targeted or as by-catch (Dulvy et al., 2014; MacNeil et al., 2020; Pacoureau et al., 2021). Although some elasmobranch fisheries can be sustainably managed (Simpfendorfer and Dulvy, 2017), the market demand for shark and ray products typically leads to overexploitation (Clarke et al., 2006; Dulvy et al., 2014).

The rapid global decline of elasmobranch populations requires collaborative management and conservation measures to ensure the long-term benefits of these populations to the wider ecosystem, including, where sustainable, for human resource use. Binding international trade consortia, such as CITES (Convention on International Trade in Endangered Species of Wild Fauna and Flora), regulate and provide the framework to restrict the international trade of species of priority conservation concern by creating species listing (CITES appendix I and II). Indeed, there has been an increasing number of elasmobranch listings in CITES Appendix II over the last decade with 38 of the 47 species regulated by CITES added at the 16^th^ (2013), 17^th^ (2016) and 18^th^ (2019) Conference of the Parties conventions (Booth et al., 2020). The number of Appendix II listings then more than tripled at the 19th Conference of the Parties (CoP19) in 2022 where parties agreed to add all remaining (54) species of requiem sharks (Carcharhinidae spp.), 6 species of hammerhead sharks, and 37 species of guitarfishes to Appendix II. Seven species of Brazilian freshwater stingrays were also adopted for Appendix II listing. The scale and pace of these listings (now 151 species) present an important implementation challenge for countries with large and diverse landings of sharks and rays, such as Indonesia.

As a result of substantial bycatch, Indonesian fisheries hold the world’s largest volume of elasmobranch landings (Fahmi and Dharmadi, 2015; FAO, 2022). This exploitation contributes to the high vulnerability rate of elasmobranch populations in Indonesian waters (Mardhiah et al., 2019), including the populations in its coral reef ecosystems (MacNeil et al., 2020). This is particularly concerning as Indonesia harbours almost a quarter of the world’s elasmobranch diversity (Ali et al., 2018; Ali et al., 2014). Despite this, export volumes of elasmobranch products from Indonesia represent only a small fraction of its landing volume (FAO, 2021), which likely reflects its communities’ high dependency on shark and ray as an alternative protein source (Dharmadi et al., 2019b; Muttaqin et al., 2018; Prasetyo et al., 2021). Several measures have been established by the Indonesian authorities to reduce the decline of elasmobranch populations, such as: increasing the number of protected species, extensive outreach programmes, improvement of data collection and stock assessment, expansion of marine protected areas, as well as the establishment of port state measures to combat illegal fishing (Booth et al., 2018; Dharmadi et al., 2015; Nugraha et al., 2020; Oktaviyani et al., 2019).

The issue around elasmobranch fisheries is rendered even more challenging by the myriad of shark and ray product derivations, which add another layer of complexity (Dent and Clarke, 2015; Safari and Hassan, 2020; Shea and To, 2017). Due to their similarity in appearance and the lack of distinctive features in most derivative products, elasmobranch species can be deliberately or accidentally mislabelled by those involved in the trade (**Figure 1**). The general lack of transparency in the trade of living resources is an ongoing concern for fisheries and conservation management (Naaum and Hanner, 2016) and can have a negative impact on stock management, and damages the reputation of entire sectors and countries (Cawthorn and Mariani, 2017; Naaum and Hanner, 2016). Furthermore, the continuous increase of elasmobranch species listed in the CITES Appendices requires constant improvements of national and transnational capabilities in monitoring the supply chain (Pavitt et al., 2021).

**Figure 1.**
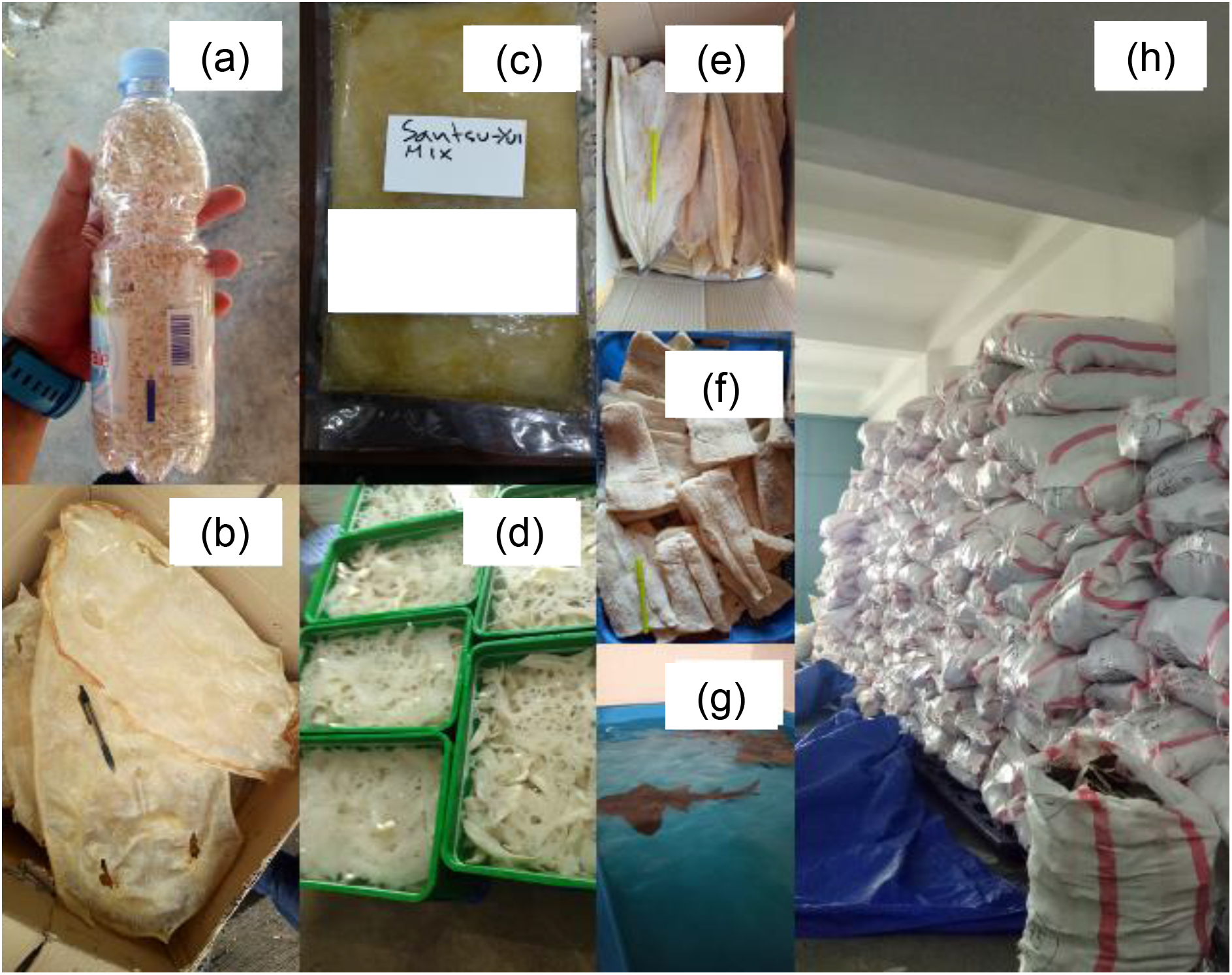
Condition of inspection and some derivative products from sharks and rays i.e. shark teeth (a); processed ray skin (b); shredded fins ‘hissit’ in brine ready for exporting to Japan (c); blue shark cartilages soaked for processing (d); dried meat from small sharks (e); dried meat from a large shark (f); live bowmouth guitarfish for the aquarium market (g); and dried fins of silky and hammerhead sharks waiting for quota to export (h).

The rapid development of DNA-based diagnostic tools offers an ever-expanding option for wildlife identification, which have greatly assisted elasmobranch biology and forensics. Established DNA barcoding (Shivji et al., 2002) and mini-barcoding (Fields et al., 2015) approaches can robustly identify species in fresh and processed samples. However, these traditional DNA barcoding methods require longer processing time and high costs for their sequencing processes. More recently, advances in real-time PCR have eliminated the sequencing stage, thereby allowing species identification to be conducted in the field. This approach uses target-specific primers and fluorescent dyes to detect the presence of the targeted nucleic acid template during PCR amplification and has been successfully applied to detect several CITES-listed shark species in a single run tube (Cardeñosa et al., 2018) and Multiplex LAMP (Lin et al., 2021). However, given their reliance on species-specific primers and probes, these methods are better suited to screening large numbers of specimens from one or few species rather than from a wide variety of species. Thus, the need remains for a fast and easy way to identify any sample, by-passing the need to design species-specific assays.

This issue is particularly glaring when inspectors are dealing with multiple types of products from different species across many locations and with a limited timeframe to investigate species compositions (Prasetyo et al., 2021). This year, the magnitude of the challenge has more than tripled, with the number of CITES-listed species going from 47 to 151 (CITES, 2022; Collyns, 2022). Since CITES regulations still allows species listed on Appendix II to be traded by considering the sustainability of exploitation through a Non-detrimental Findings (NDF) framework, trade monitoring is more crucial than ever before.

In an attempt to circumvent the limits of species-specific methods, a universal single-tube assay marketed as FASTFISH-ID^™^ was recently developed for use in the seafood industry (Naaum et al., 2021). This method uses LATE (Linear-After-The-Exponent) PCR to amplify one strand of the full 650bp COI barcoding region (Sanchez et al, 2004), and uses a set of fluorescent probes to target two distinct mini-barcode regions selected for their high intra-specific variability which will then produce unique species-specific fluorescent signatures (Naaum et al., 2021). The fluorescent signatures are then compared to those kept in a cloud-based library of verified specimen signatures.

However, this approach and its libraries were originally designed and validated for bony fishes (Naaum et al., 2021) and no elasmobranch fluorescence fingerprints are publicly available in the FASTFISH-ID^™^ cloud. We therefore chose to test i) whether the existing FASTFISH-ID^™^ diagnostics could produce a diverse range of fluorescent signatures unique and specific to each of the 28 elasmobranch species frequently found in Indonesian trade; and ii) whether a deep machine learning method could quantitatively assign signatures to the correct species, irrespective of the visual appearance of the fluorescence. Deep learning algorithms are highly flexible and well suited for undertaking these tasks (LeCun et al., 2015; Malde et al., 2019), and have recently been applied in marine science, including fish size estimation (Garcia et al., 2019), bycatch detection and shark identification from photos and videos (Jenrette et al., 2022; Peña et al., 2021; Sharma et al., 2018). Our findings indicate that this portable, universal methodology performs well even for ‘non-target’ elasmobranch species, and with further refinement, it can become a powerful tool to combat the illegal trade of endangered sharks and rays.

## Results

### Fluorescent signature of species

After filtering and removing 33 inconsistent runs, 357 pairs of fluorescent signatures from 28 species were generated, including 14 sharks and 14 rays, with 22 of those species (12 sharks, 10 rays) being CITES-listed species. Within 2.5 hours, all types of samples - from fresh to processed samples sourced from different body parts - were amplified and produced one or two fluorescent signatures (referred to as BS1 and BS2 for barcode segment one and barcode segment two) (**Tables S.1 and S.2**. These two barcode segments refer to the two mini-barcode regions within the amplified COI target sequence that emitted fluorescent to be read by the real-time PCR machine.

Many species were distinguishable using a combination of both barcode segments and had unique signatures, such as *Alopias pelagicus* (pelagic thresher), *A. superciliosus* (bigeye thresher) and *Isurus paucus* (longfin mako shark). However, some species displayed probe-barcode hybridisation difficulties (see Methods), with more shark species (7) than ray species (3) being affected, namely *Carcharhinus falciformis* (silky shark), *C. longimanus* (oceanic whitetip shark), *I. oxyrinchus* (shortfin mako shark), *Lamna nasus* (porbeagle shark), *C. brevipinna* (spinner shark), *Galeocerdo cuvier* (tiger shark), *Prionace glauca* (blue shark), *Rhynchobatus laevis* (smoothnose wedgefish), *Glaucostegus typus* (giant shovelnose ray), and *Pristis pristis* (Largetooth sawfish). Nevertheless, some of the species displaying poor probe-barcode hybridisation remained distinguishable using the alternative barcode segment **(Table 1 and Figures S.1-4**).

**Table 1.**
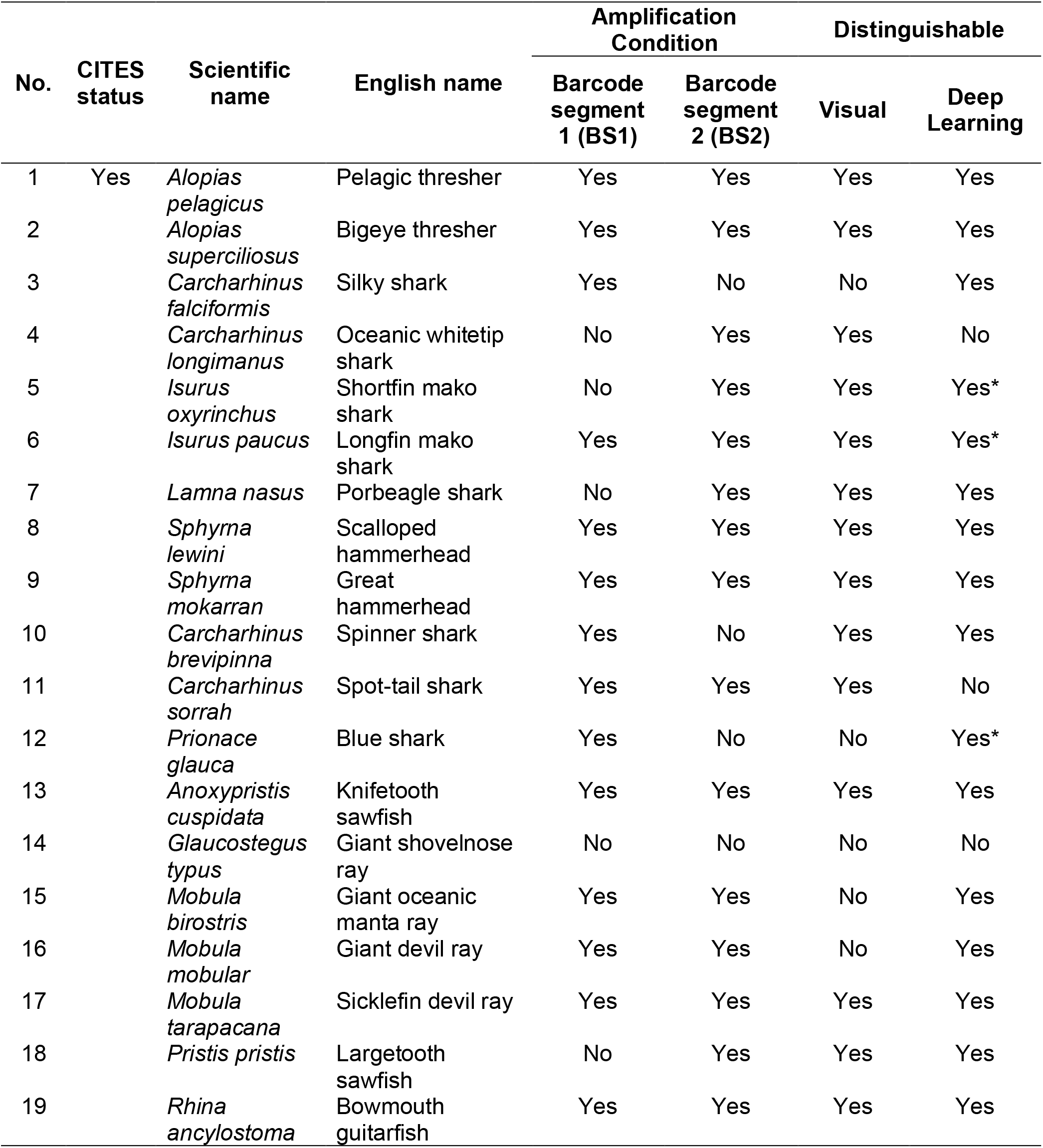
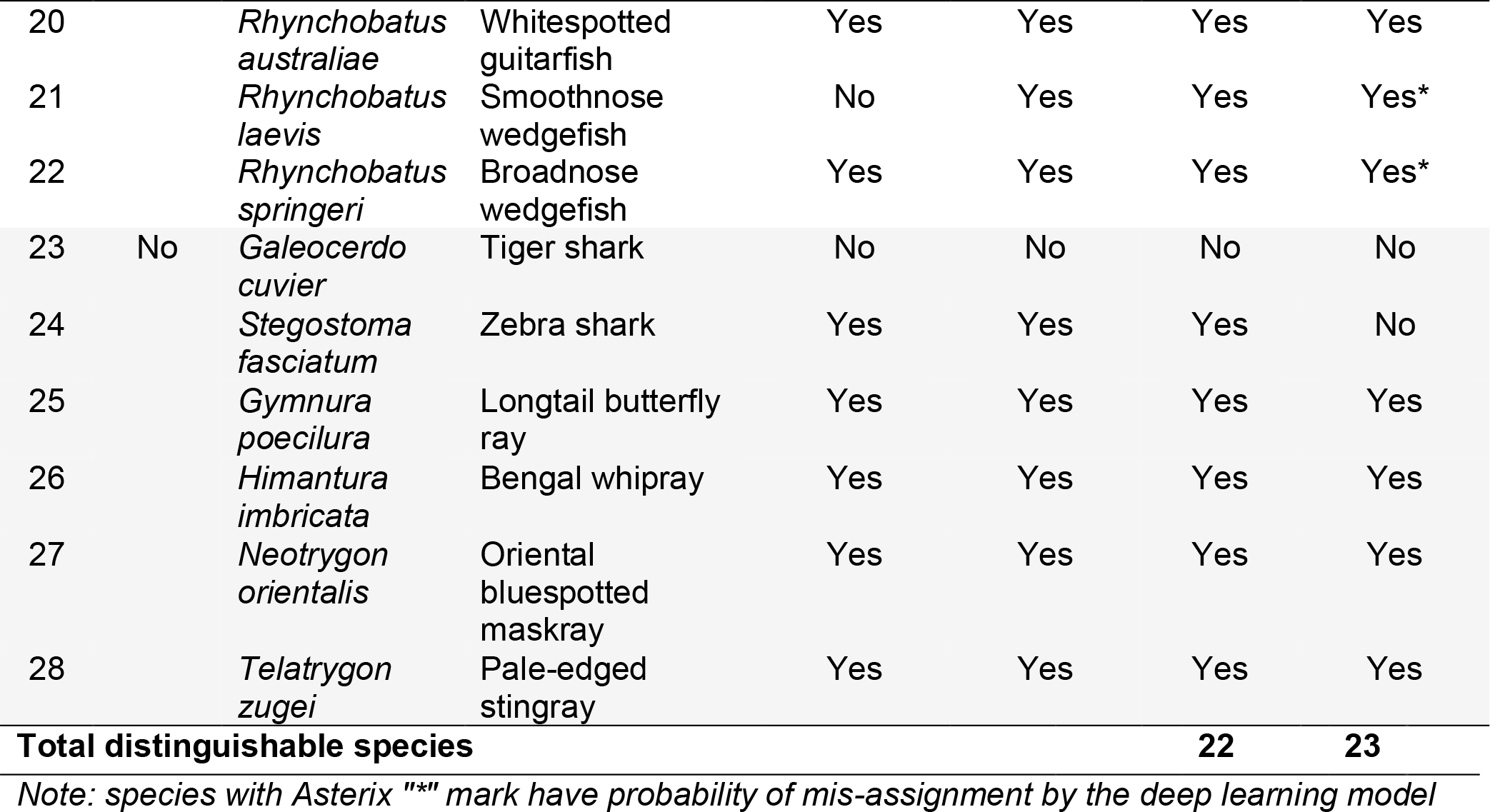
Amplification conditions of each species using the targeted segments using the FASTFISH-ID technology. Amplification condition denotes whether the species amplified at either or both segments (BS1 and BS2) and whether the species was distinguishable from all other species by its fluorescent signature(s) and deep learning.

Based on visual evaluations, the generated melt curves showed different fluorescent signatures for closely related species, such as thresher sharks (*Alopias* spp.) and hammerheads (*Sphyrna* spp.; **Figure 2**). Across the two species of thresher sharks, FASTFISH-ID^™^ produced visually distinguishable curves in BS1 at the initial stages of the hybridization process and produced a similar drop at ~74-79°C, while the signatures in BS2 were clearly distinct in the initial stages (about 42-47°C). Some species, on the other hand, have virtually identical BS1 signatures but are distinguishable using BS2, such as in the case of zebra shark (*Stegostoma fasciatum*) and spot-tail shark (*C. sorrah*) (**Figure 3**). However, there are problematic species pairs that have highly similar signatures with both segments and therefore appear visually indistinguishable. This is the case between the tiger shark and giant shovelnose ray, between the silky and blue sharks, and between the giant oceanic manta and giant devil ray (two *Mobula* species), which have nearly identical signatures in both barcode segments (**Figure 4**). Overall, six out of 28 species were deemed visually indistinguishable, four of which are CITES-listed. We also found seven species that amplified inconsistently; shortfin mako shark (*Isurus oxyrinchus*), oceanic whitetip shark (*C. longimanus*), porbeagle shark (*Lamna nasus*), tiger shark (*Galeocerdo cuvier*), largetooth sawfish (*Pristis pristis*), giant shovelnose ray (*Glaucostegus typus*) and smoothnose wedgefish (*Rhynchobatus laevis*). It was observed that the right-most trough in the BS1 fluorescent signature labelled “TM” corresponds to ThermaMark, an internal marker for correction of artefactual temperature variation (**Figure S.5**). However, in BS2, some segments were amplified and unique for each of these species.

**Figure 2.**
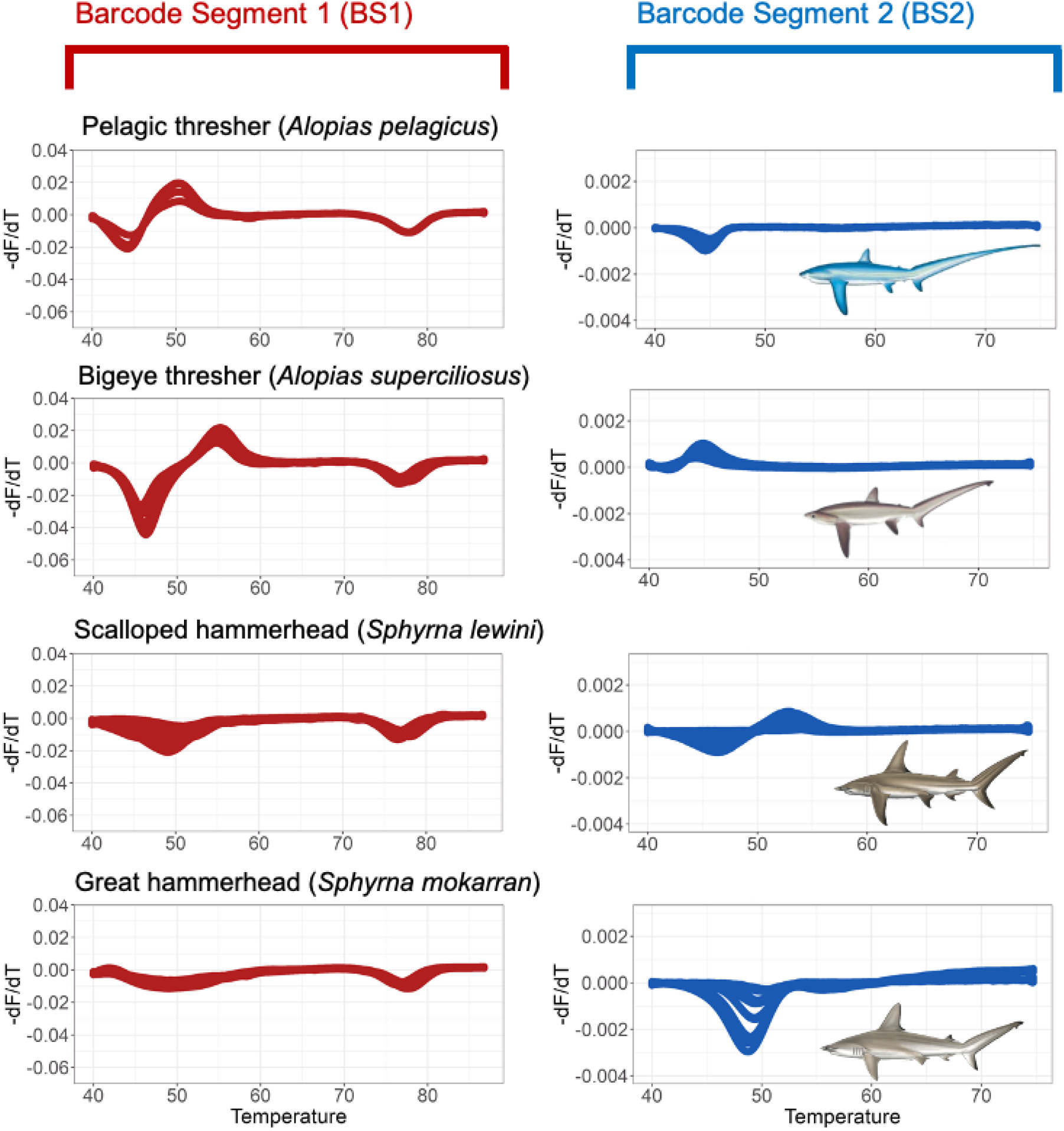
Some species that have visually distinguishable signatures in both barcode segments i.e. pelagic thresher, bigeye thresher, scalloped hammerhead and great hammerhead.

**Figure 3.**
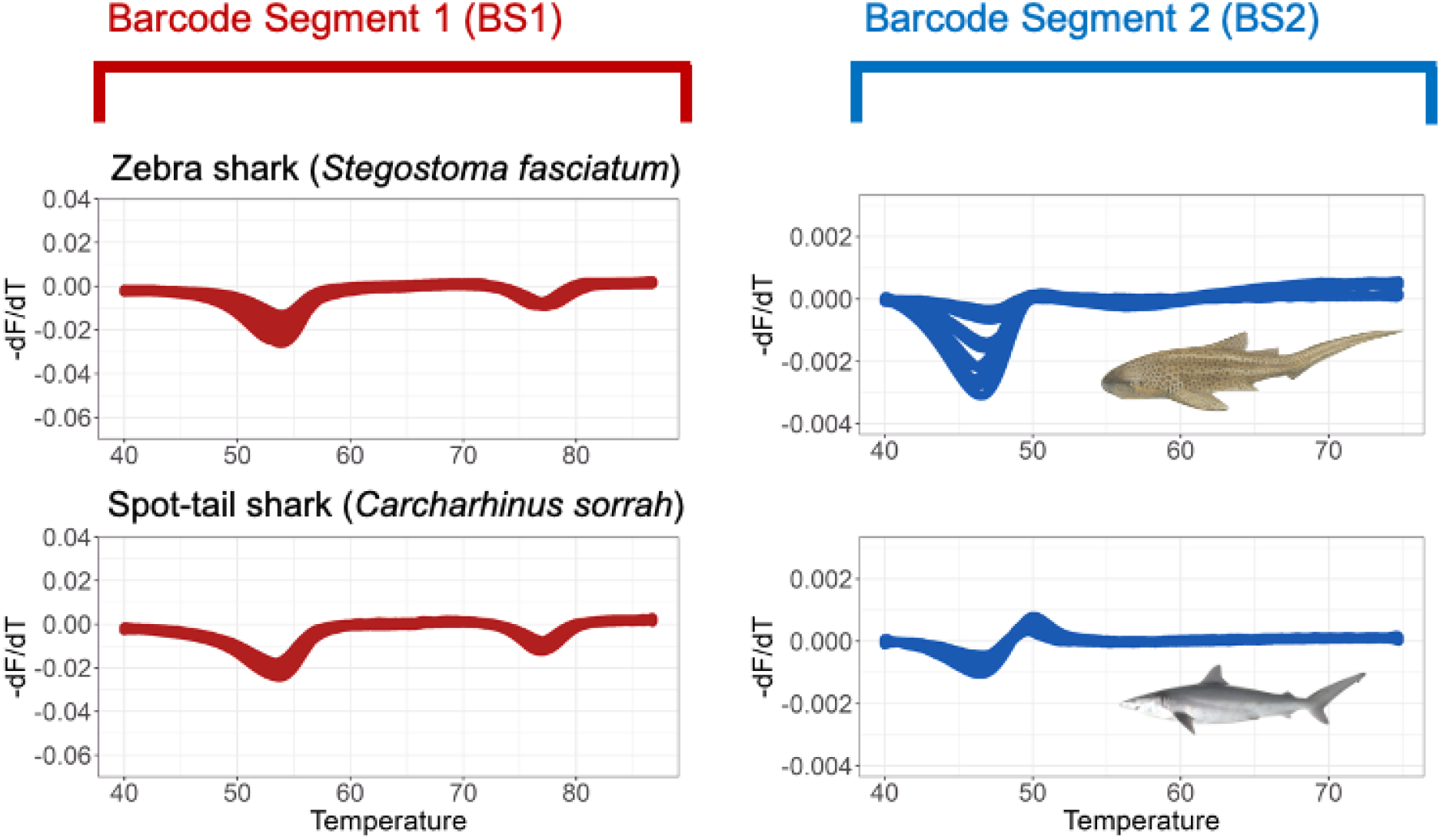
Some species that have similar signature in one barcode segment but visually unique in other segment i.e. zebra and spot-tail shark.

**Figure 4.**
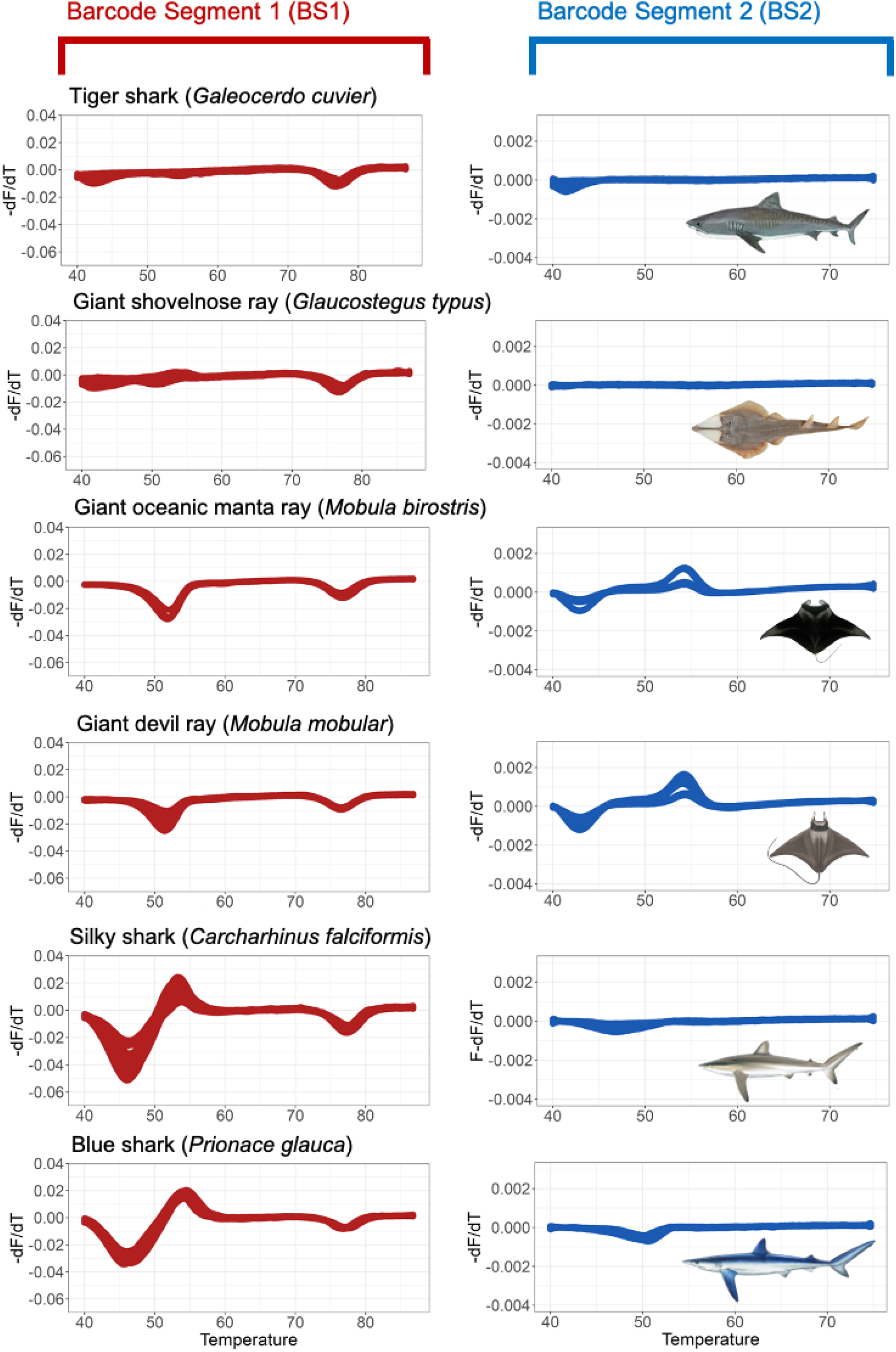
Problematic species that visually have highly similar signatures at both barcode segments i.e. tiger shark and giant shovelnose ray; giant oceanic manta ray and giant devil ray; silky shark and blue shark.

Half of the samples were highly processed products, but they still amplified well. In some of these, there were differences in the intensity of the signatures, as reflected in signature variation from BS2 of great hammerhead, zebra shark and bowmouth guitarfish (**Figure 2, 3 and S.4**), which may in part be ascribed to the actual state of degradation of the original DNA template.

### Machine learning for species assignment

We transposed data for the training sets and then used fluorescence values at 8,152 temperature intervals (>4,000 per each barcode segment) as variables and identified variable importance as a key feature for species assignment. We ranked variable states according to their relative importance, scaled importance and percentage of variance explained, for each barcode segment (see **Table S.3**). We generated 301 potential deep learning models, aiming for high accuracy and minimizing error. The best deep learning model was chosen as the one with the highest accuracy (98.20%; **Table S.4**). When the model was applied to melt curve data from the independent specimens, accuracy dropped to 79.41%, with 54 out of 68 specimens correctly assigned (**Figure 5**). Mis-assignments were consistent with the species that also proved problematic during visual assessments, i.e. the spinner and blue shark. The model also mis-identified spot-tail shark as zebra shark despite it visually having a unique signature in BS2 (**Figure 3**). During the testing, some samples from hammerhead sharks (*Sphyrna* spp.), smoothnose wedgefish (*Rhynchobatus laevis*), and broadnose wedgefish (*Rhynchobatus springeri*) were assigned to the wrong species, even though each of these species had their own unique fingerprint (**Figures S.1-4**).

**Figure 5.**
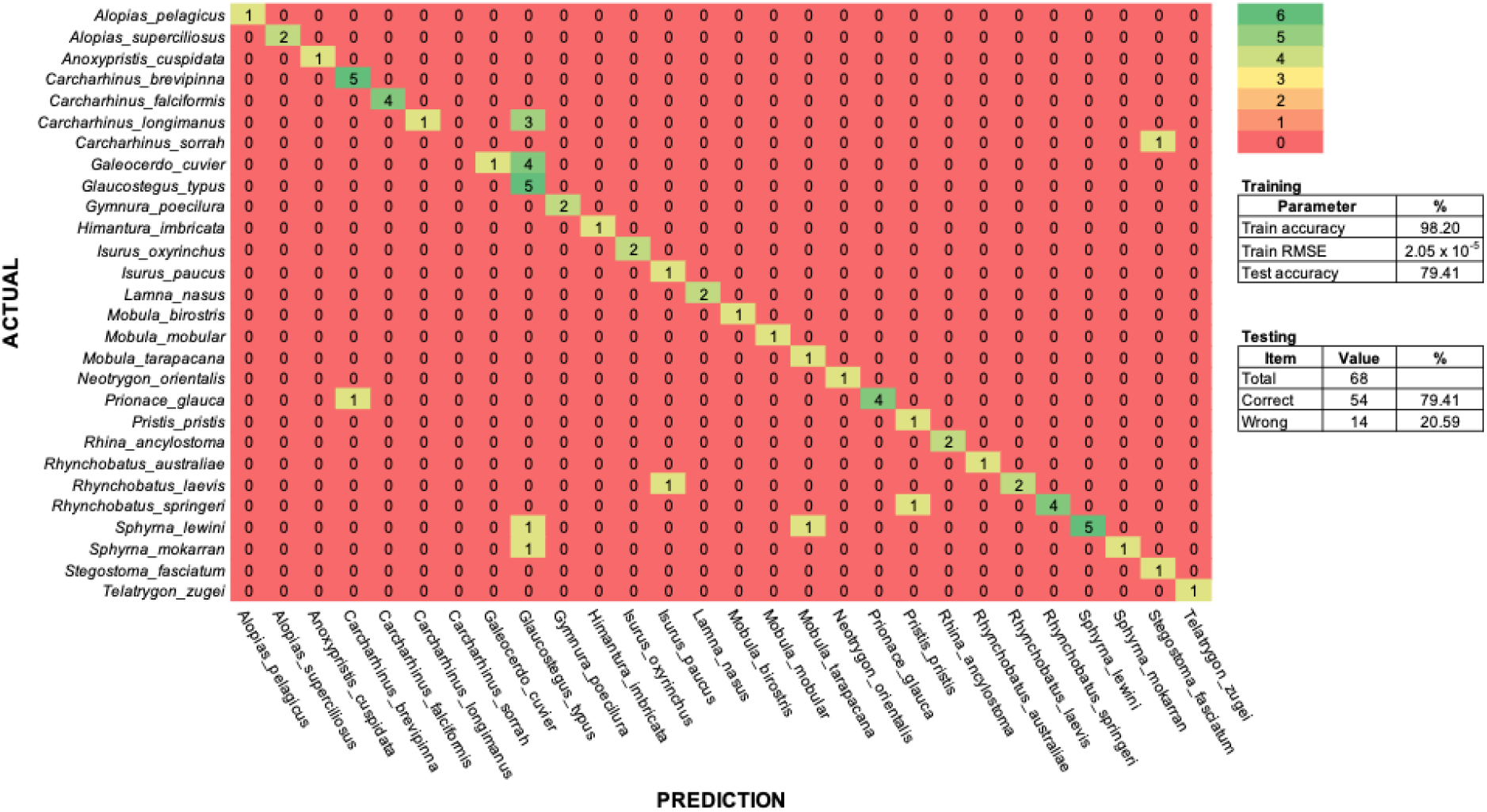
Confusion matrix of 28 shark and ray species assignment.

## Discussion

Within a couple of hours and without the need to adjust the existing FASTFISH-ID^™^ assay from teleost fish to elasmobranchs, this real-time PCR method offered a portable monitoring tool that reliably enabled the identification of 25 elasmobranch species (20 of which are CITES-listed). The device used to conduct the runs, the MIC, is a convenient portable real-time PCR thermocycler weighing no more than 2 kg and allowing for the simultaneous inspection of 48 specimens per run (Naaum et al., 2021). More importantly, the use of probes targeting mini barcodes with high inter-specific variation offers a universality that other qPCR-based assays do not currently provide, and the automatic amplification of the full COI barcode as part of the same reaction offers downstream opportunities for further in-depth screening, if necessary.

While existing genetic-based monitoring tools continue to be useful in many situations (Fields et al., 2015; Shivji et al., 2002)(Cardeñosa et al., 2018; Lin et al., 2021), FASTFISH-ID^™^ seems poised to significantly expand the horizons of DNA-based control: alongside its speed, portability, and universality, the method exhibits single nucleotide resolution (Rice et al., 2014) which can minimize the risk of similar fluorescent signatures, particularly when more species are added to a reference library (Naaum et al., 2021). This is a particularly compelling argument for its implementation, as CITES lists are likely to continue to expand in the future. Additionally, the amplification of the whole COI universal barcode segment embeds a forensic dimension (Dawnay et al., 2007) that is not necessarily afforded by other portable tools.

A difficulty typically encountered in genetic-based trade monitoring is the handling of processed products, and this is particularly true for elasmobranchs which tend to be heavily processed in a variety of ways (Dharmadi et al., 2019a; Muttaqin et al., 2018). Despite the issues of fragmented DNA due to the effect of various processing techniques (Shokralla et al., 2015), FASTFISH-ID^™^ shows notable robustness and reliability, with 83.6% of processed samples yielding reliable melt curve profiles (51 of 61 processed samples). Since FASTFISH-ID^™^ uses real-time PCR and relies on fluorescent signatures, some species display variation in signature amplitude (the variation in peak heights and valley depths) especially when the DNA was degraded, as observed with processed products and displayed by the signature of both hammerhead species on BS2 (**Figure 2**). This deviation may be problematic for species assignment, especially when the assignment depends on a deep learning algorithm. The high probability of the features being similar to those of other species caused misassignments. Other issues that may have occurred is variation in the fluorescence signature from the same species. This could be due to single nucleotide polymorphisms (SNPs) within species or possibly to contamination in the case of the BS2 signature of the pale-edged stingray (*Telatrygon zugei*; **Figure S.4**).

Visual assessment could distinguish 22 species out of 28 with more than half of these (N=17) being CITES-listed. Even in this preliminary phase, the method could therefore readily be applied by inspectors–without the application of computational tools – and reliably reveal cases of illegal activities. Three pairs of species had spectral features that are difficult to distinguish, e.g. these ambiguities were present between tiger shark and giant shovelnose ray, between two species of *Mobula* rays (giant oceanic manta ray and giant devil ray), and between silky and blue shark **(Table 1 - Visual**). Thus, it must be acknowledged that the barcode segments have the same sequence of nucleotides and produced similar signatures for those species. The technology was originally designed for bony fish (Naaum et al., 2021), and the database is currently being expanded to various important species that are globally traded as seafood. Yet, the much lower diversity of elasmobranchs (~1/30^th^ that of teleosts) will make any effort to produce spectral reference databases a far less onerous task than that currently encountered with bony fishes. Whilst it has been known that the COI gene is more slowly evolving in chondrichthyans than teleosts (Moore et al., 2011; Naylor et al., 2012), this is seldom a major issue in most DNA barcoding applications (Fields et al., 2018; Griffiths et al., 2013; Hobbs et al., 2019), so an optimised iteration of the FASTFISH-ID^™^ method is poised to be transformational for elasmobranch conservation and management. A qualitative investigation on the full length of COI sequences (Sanger sequencing results) based on visual and simple comparison (https://www.bioinformatics.org/sms2/ident_sim.html) revealed that for those problematic three pairs of species mentioned above for that particular segment, there is a high degree of similarity in their sequence (70-98%), although this seems unlikely as the method is extremely sensitive and easily distinguishes between sequences that differ by a single nucleotide (Sirianni et al., 2016).

In the absence of an online reference database of elasmobranch fluorescent signatures, machine learning was developed for this study. One of the machine learning applications is pattern recognition (Jenrette et al., 2022; Trentin et al., 2018). Deep learning (also known as deep structured learning) is broadly applied in machine learning applications, especially pattern recognition (Jenrette et al., 2022; Trentin et al., 2018) and has advantages in its flexibility to develop learning styles i.e. supervised, semi-supervised or unsupervised (LeCun et al., 2015; Malde et al., 2019). Deep learning models have been chosen and deployed with independent testing datasets to measure their accuracy. We found that the accuracy of our test model was 79.41%, which is lower than the training accuracy (98.20%; **Table S.5**), and yet the model could identify similar species that could not be distinguished visually. In fact, the model enabled us to differentiate the two *Mobula* species that have similar signatures in both barcode segments. Machine learning could also recognize silky shark, a problematic species for the authorities as the species belongs to the Carcharhinidae, a diverse family that has plenty of look-alike species. In particular, the silky shark spectral profiles appeared visually indistinguishable from blue shark. However, the new CITES listing agreed during CoP19 added all requiem sharks into Appendix II (including blue shark along with the other 53 species shark from Carcharhinidae family) will make implementing action manageable since requiem sharks make up a large proportion of the products found in the global shark fin trade hubs in China (Cardeñosa et al., 2022). Although international trade in all requiem sharks will now be regulated, a Non-Detriment Finding (NDF; CITES’s mechanism that allows certain species listed in Appendix II to be traded with strict quotas) which is specific to each species will still require the capability of identification at the species level.

Five out of 28 species could not be assigned accurately using the model, i.e. between spot-tail and zebra shark as well as mis-assignments among oceanic whitetip shark, tiger shark and giant shovelnose ray **(Table 1 – Deep Learning**). Curiously, there were also mis-assignments for species that had quite unique fluorescent signatures. We argue that these mis-assignments could be due to variation in amplitude, where some species actually have similar signatures, but different amplitudes (Cusa, 2021) the cause of which is undetermined, but could be due to degraded DNA. For instance, the signature in BS2 of zebra shark has high amplitude variations that may challenge the model to assign the species (**Figure 3**). Increasing training datasets may be required as this should improve the robustness of the model (LeCun et al., 2015), while future re-tailoring of the barcode regions to elasmobranch variation may also remove some of the within-species noise. Despite the assignment problems, when we combine visual and deep learning assignments, we could distinguish 25 out of 28 species, 20 of which are listed in CITES Appendix II.

## Limitations of the study

The probe hybridization problems (which occurred when the barcode segments have a high degree of mismatches with the designed probes) encountered in seven species prevented the machine learning tool from adequately assigning fluorescent signatures to a given species. Since BS1 failed to hybridize for most of these species, the species assignment in these cases was solely reliant on BS2, which, in many cases also exhibited poor hybridization. To address this issue, it seems that going forward the designing of new probes tailored to elasmobranch sequence variation will be a necessary solution to increase the versatility and reliability of FASTFISH-ID^™^. An increased set of elasmobranch species may also inflate mis-assignments due to the higher degree of similarity among species in both visual-based or machine learning-based systems. There is also limitations in using fully supervised deep learning approaches in the selection of important features from highly variable training sets (e.g. signatures from the two barcode segments) (Hantak et al., 2022). The addition of more species to the database will require more training images. However, with such improvements, this method will help authorities (i.e. fish inspectors, customs and quarantine officers) by providing a single, agile testing option, at any point in the supply chain, to disentangle the complexity of the shark and ray product trade, and ultimately reduce the consequential risk of extinction for these endangered and iconic taxa.

## Acknowledgements

We thank all collaborators of the project Building Capacity to reduce illegal trading of shark products in Indonesia, funded under the Illegal Wildlife Trade (IWT) Challenge Fund number IWT057 and the University of Salford R&E strategy funding, the Ministry for Marine Affairs and Fisheries (MMAF) – Republic of Indonesia, the University of Salford (UoS), the Centre for Environment, Fisheries and Aquaculture Science (Cefas) and the Rekam Nusantara Foundation – Indonesian (REKAM). We also thank the officers and staff of B/LPSPLs (‘Balai/Loka Pengelolaan Sumber Daya Pesisir dan Laut’; Institute for Coastal and Marine Resource Management) and Fish Quarantine for helping during field work. We thank staff of the Wildlife Conservation Society – Indonesian Program (WCS-IP) for their support during project initiation. In addition, A.P.P. would personally like to thank Mr. Dharmadi and Dr. Hawis Maddupa for their legacy and supporting the younger generation of scientists seeking to make an impact in conserving elasmobranchs in Indonesia. We also thank Marc Dando for the use of his scientific illustrations of elasmobranchs.

## Author contributions

Conceptualization (APP, ADM and SM), Funding acquisition (SM and JMM), Methodology (APP, MC, ADM and SM), Resources (ADM and SM), Investigation (APP and MC), Formal Analysis and Visualization (APP), Project Administration (APP, JMM, FA, ADM and SM), Supervision (ADM and SM), Writing—original draft (APP, ADM and SM), Writing—review & editing (MC, JMM, FA and EM).

## STAR Methods

### Resource availability

#### Lead contact

Further information and requests for resources and reagents should be directed to and will be fulfilled by the lead contact, Andhika Prasetyo or Allan McDevitt.

#### Materials availability

This study did not generate new unique reagents. FASTFISH-ID^™^ reagents were manufactured by Ecologenix, LLC. Natick, MA - USA.

#### Data and code availability

- Data is archived at the Google Drive and are publicly available of the date of publication database: https://bit.ly/FASTFISH-ID_MS_Supp_Datasets.
- All original code is deposited at the Github repository and are publicly available of the date of publication database: https://github.com/andhikaprima/FastSharkID.
- Any additional information required to reanalyse the data reported in this paper is available from the lead contact upon request.

### Experimental model and subject details

Tissue sample of shark and ray specimens were collected in several sites nested in six locations across cities on Java Island, the most populous island in Indonesia (**Figure S.6**, namely Jakarta, Indramayu, Tegal, Cilacap, Surabaya and Banyuwangi. Collected specimens were gathered without prior knowledge of their exact harvest location and were available for collection at a variety of sites, such as fishing ports (FP), traditional markets (TM), processing plants (PP), export hubs (EH) and an inspector station (AU).

Sample collection was granted by research permit no.251/BRSDM/II/2020 issued by Agency for Marine and Fisheries Research and Human Resources AMFRAD, the Ministry of Marine Affairs and Fisheries (MMAF), Republic of Indonesia. Research ethics no. STR1819-45 issued by the Science and Technology Research Ethics Panel, University of Salford. Export permits no. 00135/SAJI/LN/PRL/IX/2021 (CITES-listed specimens) and 127/LPSPL.2/PRL.430/X/2021 (non-CITES-listed specimens) were granted under the authority of the Ministry of Marine Affairs and Fisheries (MMAF), Republic of Indonesia. Sample were imported into the UK under import permit no. 609191/01-42 from the Animal and Plant Health Agency (APHA), United Kingdom.

### Method details

#### Sample collection and DNA extraction

579 specimens were opportunistically collected at the above-mentioned sites and processing factories throughout January and February 2020. The tissue, which could either be fresh, frozen, partially or heavily processed, was then stored in 2.0mL screw-cap microcentrifuge tubes, submerged in 90% ethanol and stored at 4°C. DNA was extracted from samples following the Mu-DNA protocol for tissue samples (Sellers et al., 2018) with an overnight incubation at 55°C on the thermomixer with a medium mixing frequency and a final elution volume of 100 μl. All surfaces were sterilised with 50% bleach and then washed with 70% ethanol, in-between and after extracting each sample, to reduce cross-contamination risks (**Figure S.7a-b)**.

Of these, we excluded specimens of unclear taxonomy, and all species represented by less than 3 individuals. We refined the collection to 130 tissue samples (specimens) belonging to 28 species; for each species, we used three replicates per specimen as training sets (390 runs) (**Table S.1)**. We also had another 68 tissue samples without replication and used them as testing datasets (**Table S.2)**. As sampling was conducted opportunistically, we did not have an equal number of samples per species. Some species had a limited number of specimens, so we took out some training sets to be used as testing datasets. Datasets were then filtered, and ambiguous qPCR runs (i.e. poor probe-barcode hybridisation or inconsistent fluorescent signature) were removed. A poor probe-barcode hybridisation was checked using a reference point created by ThermaMark™ (TM) in the signature produced from BS1. If only ThermaMark™ (TM) amplified in the BS1 fluorescent signature, those runs would have failed to hybridize. Inconsistent fluorescent signatures within a replication or species were re-run a second time. If the re-runs kept failing, those runs were removed. In the end, we used 357 (number of replications varied by specimens) and 68 runs for training and testing datasets, respectively.

#### FASTFISH-ID^™^ closed-tube barcoding protocol

##### PCR reaction and amplification conditions

In the first instance, the FASTFISH-ID^™^ method requires the amplification of the full cytochrome c oxidase I (COI) gene (~650 bp) and in the second instance, it targets the two mini-barcodes (~80 bp) using a set of probes. PCR master mixes were prepared in low-adhesion Eppendorf tubes (Naaum et al., 2021). The major components of this method are ThermaStop^™^, ThermaMark^™^ and FASTFISH-ID^™^ Probe Mix (Ecologenix, LLC.). ThermaStop™ is a novel hot-start reagent that prevents non-specific amplification prior to the start of the reaction, while ThermaMark™ (hereafter referred as TM) is a temperature-dependent marker for correction of melt-curve analysis (Ecologenix, LLC.). The FASTFISH-ID^™^ probe mix consisted of two sets of positive/negative probe pairs labelled in two different colours that hybridize along the length of two mini-barcode regions within the amplified COI target sequence, hereafter referred to as Barcoding Segment 1 (BS1) and Barcoding Segment 2 (BS2). A M13 primer was used as a priming site that facilitates the sequencing process for eventual species validation through Sanger sequencing.

FASTFISH-ID^™^ uses asymmetric PCR to produce more single stranded amplicons which allow the probes to hybridize more easily (Sanchez et al., 2004). After amplification, mismatch tolerant positive/negative probe pairs bind to their single-stranded DNA targets. Each positive-probe is formed of a target binding sequence that is 20–35 nucleotides long and has a higher fluorescent signal when it is bound to its target sequence but a low background fluorescence when it is not. Negative-probes are only quenchers that reduce the fluorescent signal when they are bound next to their paired positive-probe. Positive/negative probe pairs can bind to both perfectly matching strands and target sequence variants with one or more nucleotide polymorphisms. This means that they can tolerate mismatches, which is one of the most important features of this technology as a single set of reagents can be used to identify a large number of species (Naaum et al., 2021). Target sequences that are similar but different, even if only by one nucleotide, almost always have different fluorescent signatures. Positive/negative probe sets therefore have the potential to discriminate among thousands of fish species and their variants (Naaum et al., 2021).

PCR amplification was performed on a Magnetic Induction Cycler (MIC) which is a real-time PCR thermocycler designed by Bio Molecular Systems™ (Upper Coomera, Queensland, Australia). Thermocycling conditions were 94°C for 2 mins, 5 cycles of 94°C for 5 secs, 55°C for 20 secs, 72°C for 45 secs, then 65 cycles of 94°C for 5 secs, 70°C for 45 secs (in total: 2 hrs, 20 mins and 44 secs). Following a total of 70 amplification cycles, the reaction leads to a 10-to 20-fold excess of single-stranded DNA which is critical for probe/target hybridization in a single closed tube (Pierce et al., 2005; Sanchez et al., 2004). At the completion of PCR, the temperature was decreased down to 40°C for 10 mins to enable the fluorescent probes in the FASTFISH-ID™ probe mix to hybridize to the excess single-stranded DNA. This step was followed by a melting curve analysis where the temperature was gradually increased from 40°C to 87°C at 0.1°C /secs with sequential fluorescent acquisition first in the MIC PCR Cycler’s Orange Channel (suitable for detection of CalRed 610-labelled probes; max excitation: 590 nm; max emission 610 nm) and then detection in the Red Channel (suitable for detection of Quasar 670-labelled probes; max excitation: 647 nm; max emission 670 nm). The first derivative of the melt curve was then used as the fluorescent signature. Species assignment was revealed by comparing a distinct mix of Cal-Red 610 and Quasar 670 fluorescent signatures (**Figure S.7c-f**). Those multiple combinations allow FASTFISH-ID™ to identify a large number of species with the same reagents (Naaum et al., 2021; Rice et al., 2012; Sirianni et al., 2016).

##### DNA barcoding and species validation

The same single strand DNA products used to generate a fluorescent signature can also be sequenced by DNA barcoding for further investigation. The sequencing protocol uses the M13 tail sequence in the FASTFISH-ID™ FISH COI HBCts excess primer (5’ CACGACGTTGTAAAACGAC 3’, a modified version of the M13F primer) as a sequencing primer to generate the sequence of the excess primer strand. By design, the excess primer-strand sequence can be queried directly in the NCBI nucleotide database (NCBI, 1988) or the Barcode of Life Database (Ratnasingham and Hebert, 2007) for species identification. In addition, we also used Fish F2 (5’ TCGACTAATCATAAAGATATCGGCAC 3’) and Fish R2 (5’ ACTTCAGGGTGACCGAAGAATCAGAA 3’) primer sets (Ward et al., 2005) for several initial specimens for comparison with HBCts excess primer (M13). Sequencing was outsourced to Macrogen Europe™. Samples were prepared according to the service provider protocols (https://www.macrogen-europe.com/services/sanger-sequencing). We also added species and/or specimens after identification using a highly degenerated primer set using a high throughput barcoding (HTB) method (A.P. Prasetyo et al., *unpublished data*); Leray-XT primer sets (313 bp). This set included the primers jgHCO2198 (5’ TAIACYTCIGGRTGICCRAARAAYCA 3’) and mlCOIintF-XT (5’ GGWACWRGWTGRACWITITAYCCYCC 3’) (Wangensteen et al., 2018).

### Quantification and statistical analysis

#### Machine learning for species assignment

Since the two probing barcode segments and the algorithm were developed for teleost fishes, they are not expected to maximise differentiation among the melt curves of elasmobranch species. Furthermore, the existing cloud-based reference library does not contain any elasmobranch signatures. We therefore developed our own species identification system by using machine learning using the H2O platform (**Figure S.7h-g)**. H2O is an open source, fast and scalable machine learning and predictive analytics platform that allows building machine learning models on big data, and improving reproducibility (Candel et al., 2016). The deep learning algorithm was deployed to address the problem of species assignment by considering its capability to arrange multiple nonlinear transformations to model high-level abstractions in data. H2O’s Deep Learning is based on a multi-layer feedforward artificial neural network (FANN) that is trained with a stochastic gradient descent using a backpropagation environment (Candel et al., 2016). Deep learning is also advantaged by extracting the optimal input representation from raw data without user intervention (Avci et al., 2021).

The fluorescent signature datasets (BS1 and BS2) were extracted, with the species identity serving as the “response”, and the transposed PCR profile temperature values being used as the predictor “variables” (each barcode fragment is recorded at about 4,000 temperature values), and fluorescent values serving as the “feature”. In deep learning, “response” refers to the individual value that served as the output (species name in our case); while “variable” refers to properties of the “response” and is evaluated through the “feature”.

The performance of deep learning algorithms depends heavily on the extracted features, so it’s important to choose the right group of features that best represent the input data (Pouyanfar et al., 2018). Data filtering was conducted to exclude poor probe-barcode hybridisation or inconsistent fluorescent signature datasets and provided the best representative of the data input. Two datasets (BS1 and BS2) were then merged by specimen ID with species name used as an input to the model. Our model was divided using a 70–30 ratio of training data to validation data (i.e. 246 and 111 runs respectively) and then tested with 68 independent datasets. Default parameters of H2O’s Deep Learning were optimized, with a process called “grid-search”, this process tried to adjust several parameters to find the optimal “stopping criteria” (list of parameters provided on **Table S.6**). We setup a “stopping criteria” to limit the computational load in searching for the best deep learning algorithm, which was based on random discreteness, the number of generated models, and model runtime (**Table S.7**). The best model was chosen based on model accuracy and Root Mean Square Error (RMSE) optimization. A confusion matrix is used to visualize model accuracy.

As for other algorithms, larger databases are required to improve predictive abilities by optimizing distributed representation, activation function non-linearity, and flexible architecture depth in terms of hidden layers and nodes (Calzolari and Liu, 2021). The main challenges in applying deep learning is overfitting due to a dominant influence on the generalization ability of a deep neural network model (Li et al., 2019). However, regularization methods such as Ivakhnenko’s unit pruning (Ivakhnenko, 1971) or sparsity (l1-regularization) or weight decay (l2-regularization) can be applied during training to combat overfitting (Bengio et al., 2013). The sparsity and weight decay were used in this study.

### Key resources table

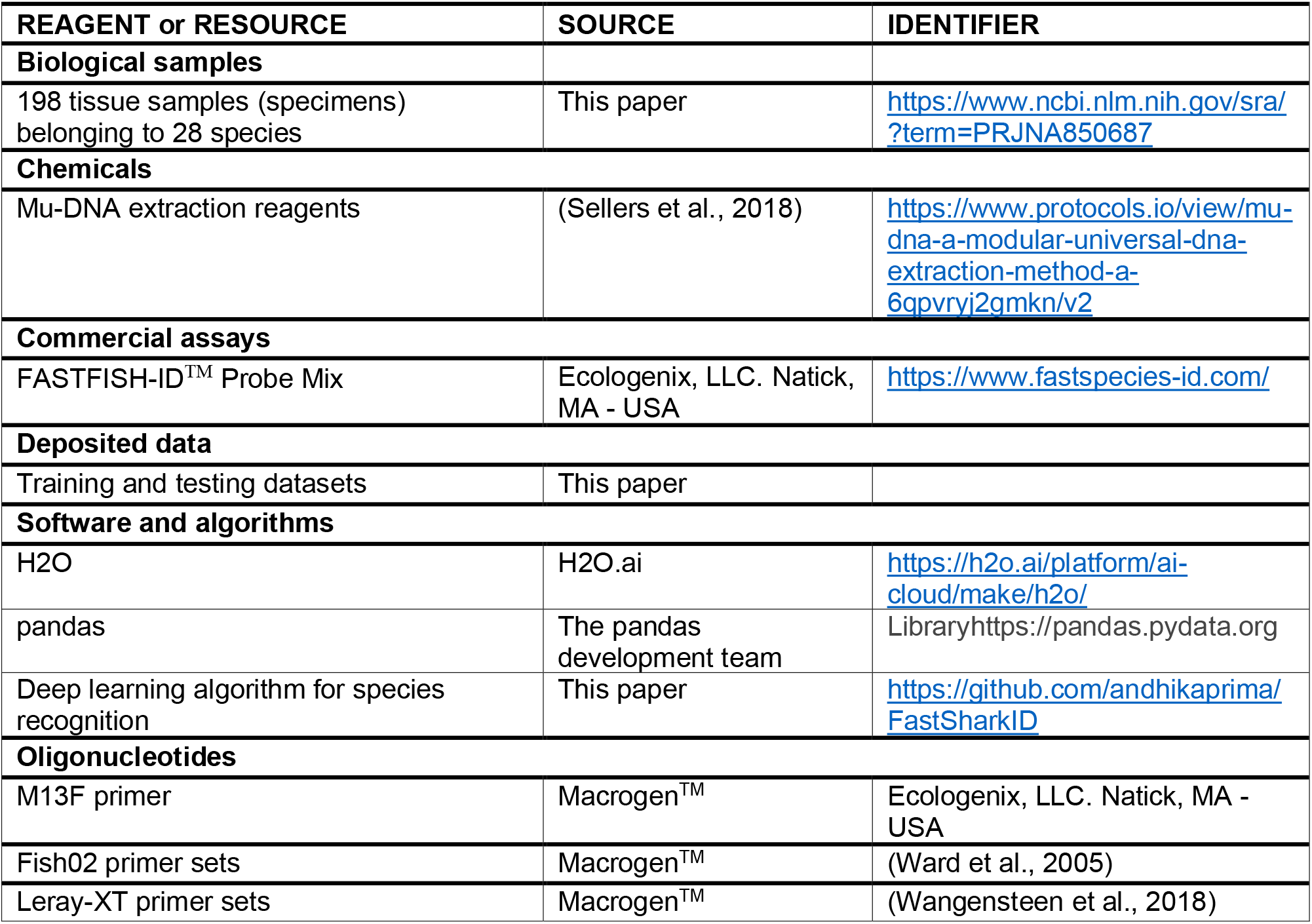

