## Supplement materials for "Universal closed-tube barcoding for monitoring the shark and ray trade in megadiverse conservation hotspots"

This file includes: Figures S1–S7 and Tables S1-S7

### Supplementary figures

**Figure S.1**. The fluorescent signatures in BS1 of 14 shark species.

**Figure S.2**. The fluorescent signatures in BS2 of 14 shark species.

**Figure S.3**. The fluorescent signatures in BS1 of 14 ray species.

**Figure S.4**. The fluorescent signatures in BS2 of 14 ray species.

**Figure S.5**. Some species which have a hybridization problem in the BS1 region. Those species only have "TM" signature (the right-most valley in the BS1, labelled with a green color), TM corresponds to ThermaMark$^{TM}$, an internal marker for correction of artefactual temperature variation.

**Figure S.6**. Sampling locations across Java Island, Indonesia. Locations are labelled with long and short codes.

**Figure S.7**. A schematic description of the stages of this study which include (a) sample collection and preservation, (b) DNA extraction of tissue samples, (c-e) sample processing using the FASTFISH-ID workflow, (f) visualisation of the RT-PCR outputs and (g and h) species classification using deep learning.

### Supplementary tables

**Table S.1**. Sample details used on the training datasets including Condition (processed/fresh), Part (of the animal), Species, ID (number), no. of replications and Sequencing technology used to identify the species.

**Table S.2**. Sample details used on the testing datasets including Condition (processed/fresh), Part (of the animal), Species, ID (number), no. of replications and Sequencing technology used to identify the species.

**Table S.3**. Variable importance in recognizing fluorescent signatures of species

**Table S.4**. Result of grid search in finding the best deep learning model

**Table S.5**. Assignment scoring of 28 species of sharks and rays

**Table S.6**. Initial value of hyper-parameters in searching for the best deep learning model using grid search method

**Table S.7**. Stopping criteria in searching the best deep learning model

### Supplementary figures

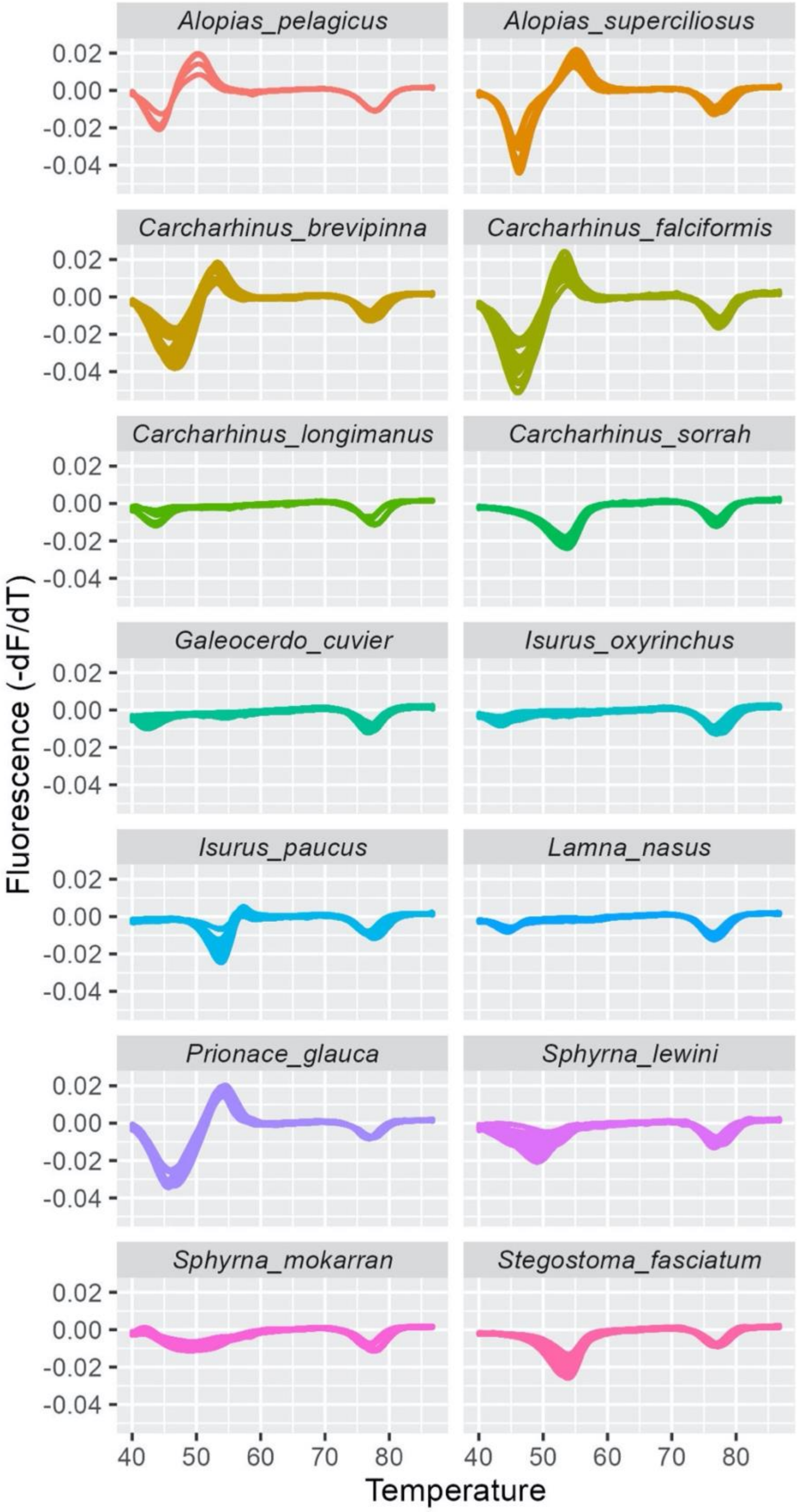


**Figure S.1**. The fluorescent signatures in BS1 of 14 shark species.

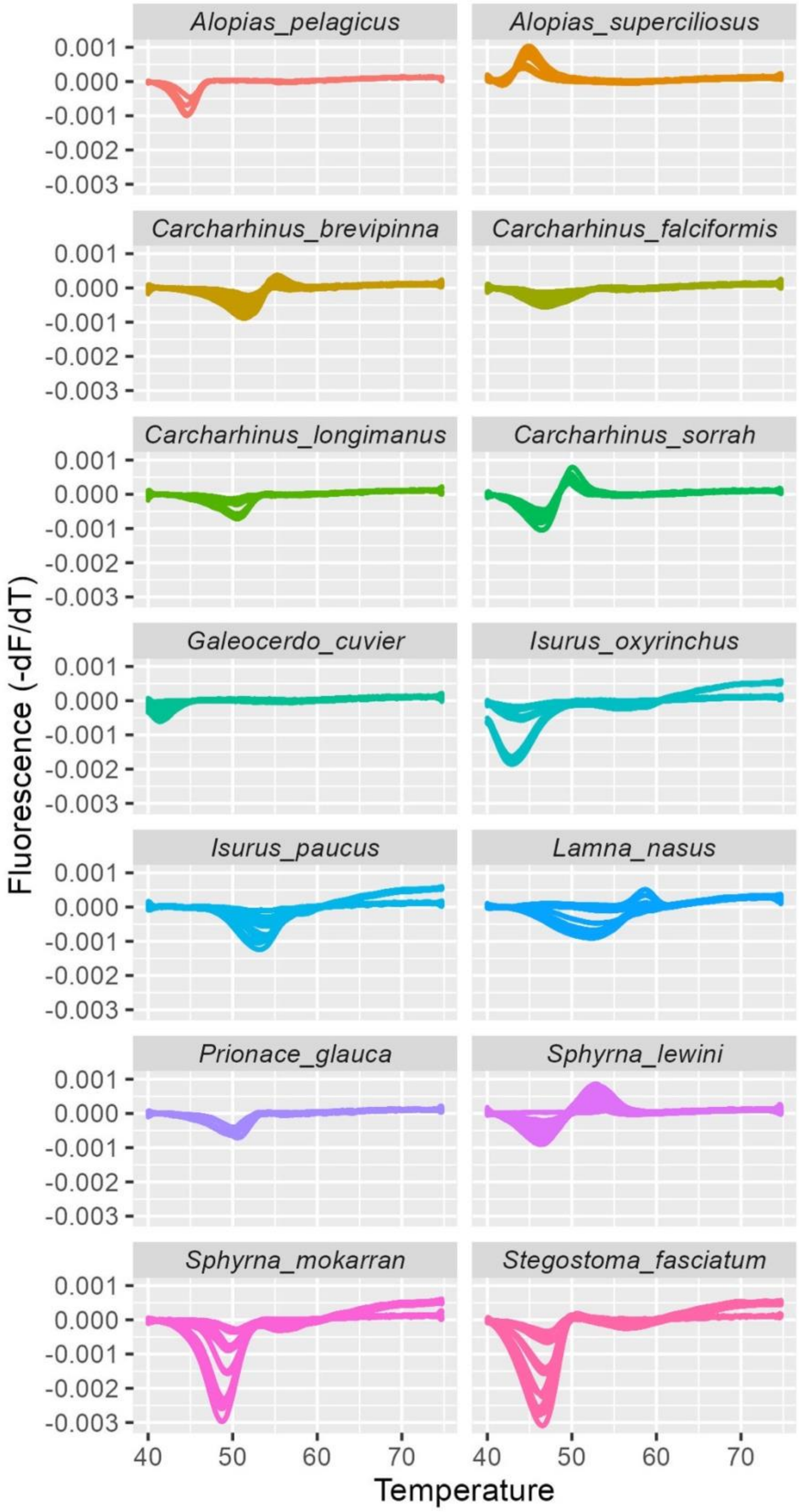


**Figure S.2**. The fluorescent signatures in BS2 of 14 shark species.

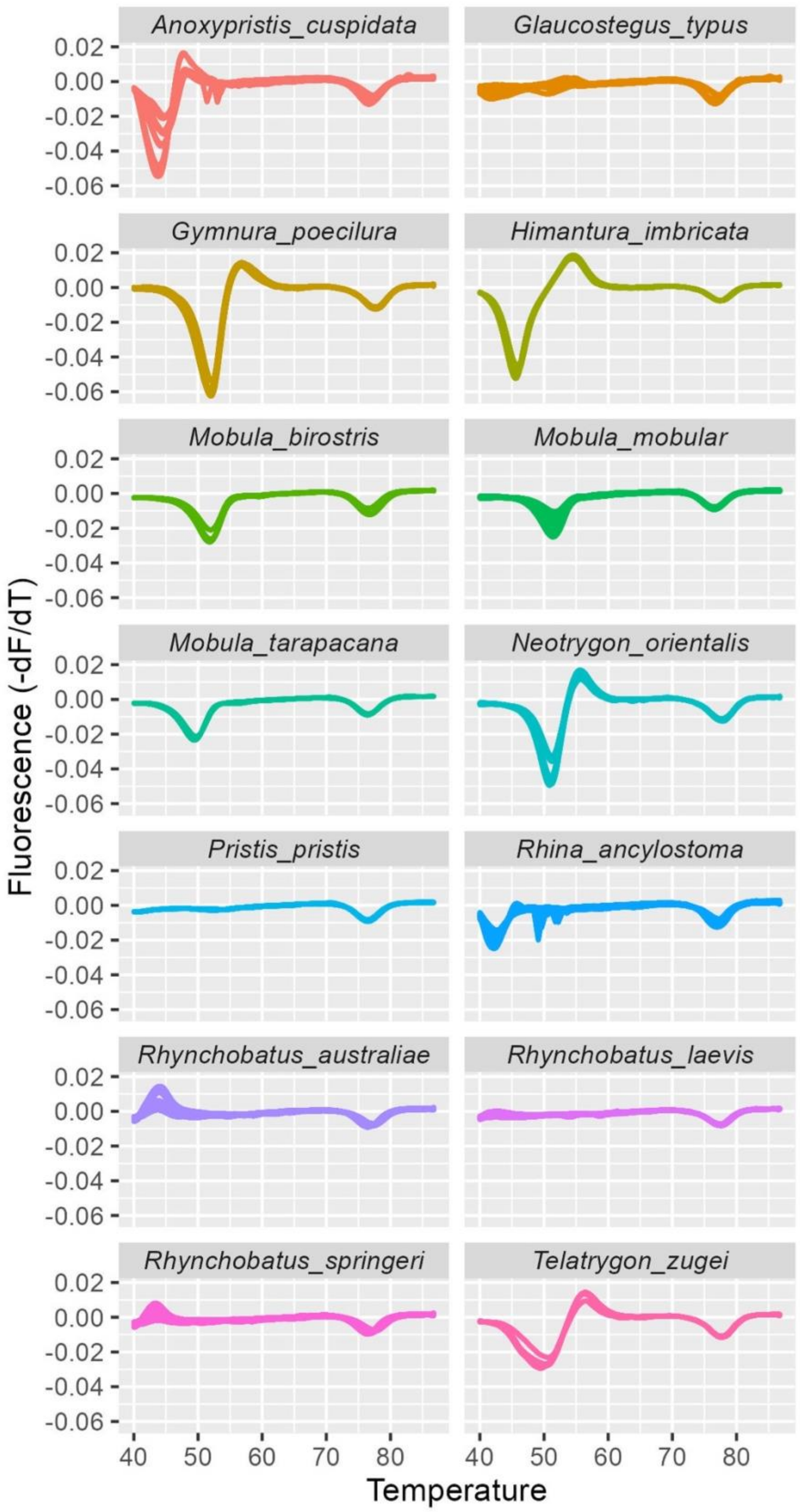


**Figure S.3**. The fluorescent signatures in BS1 of 14 ray species.

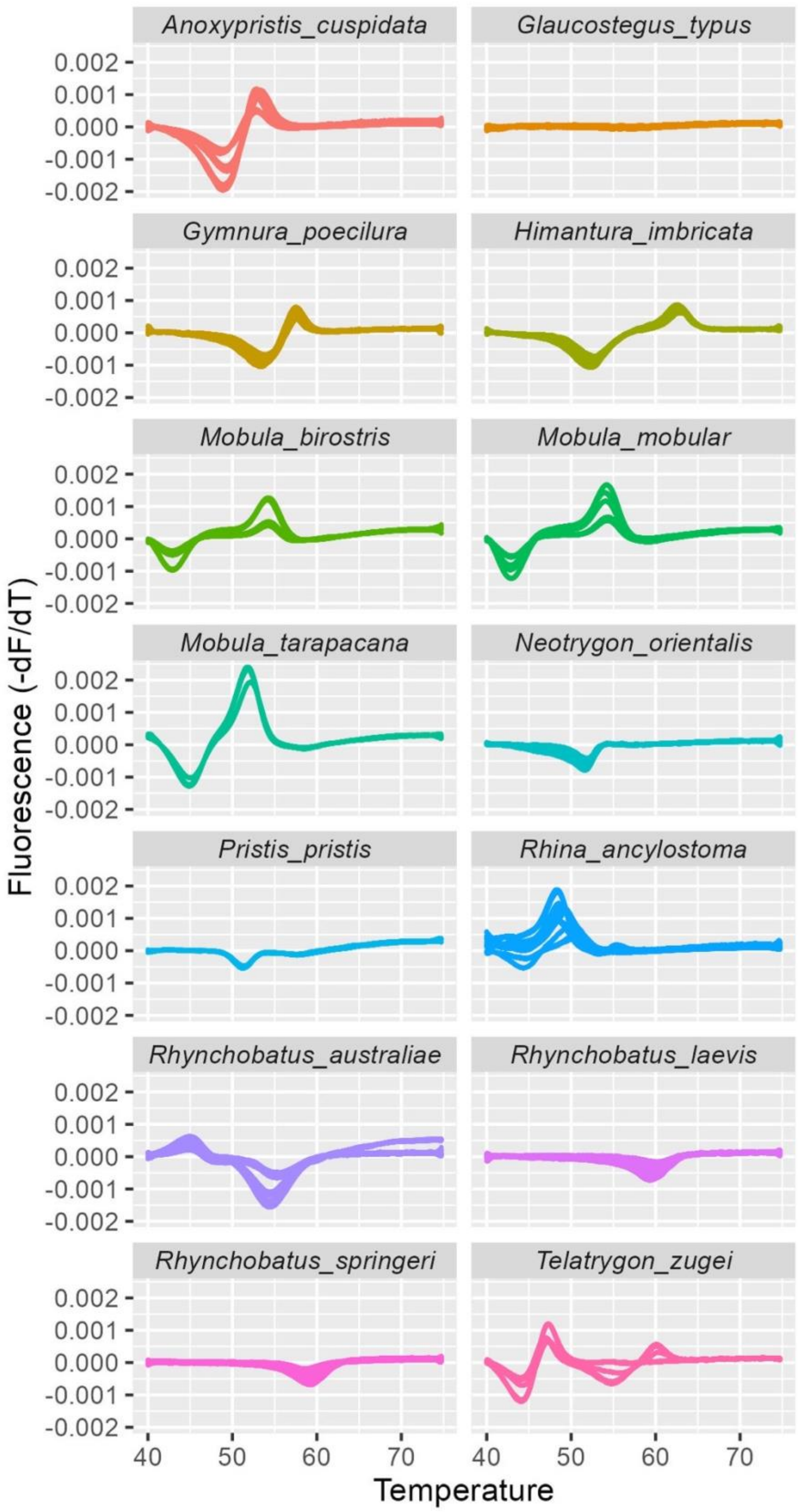


**Figure S.4**. The fluorescent signatures in BS2 of 14 ray species.

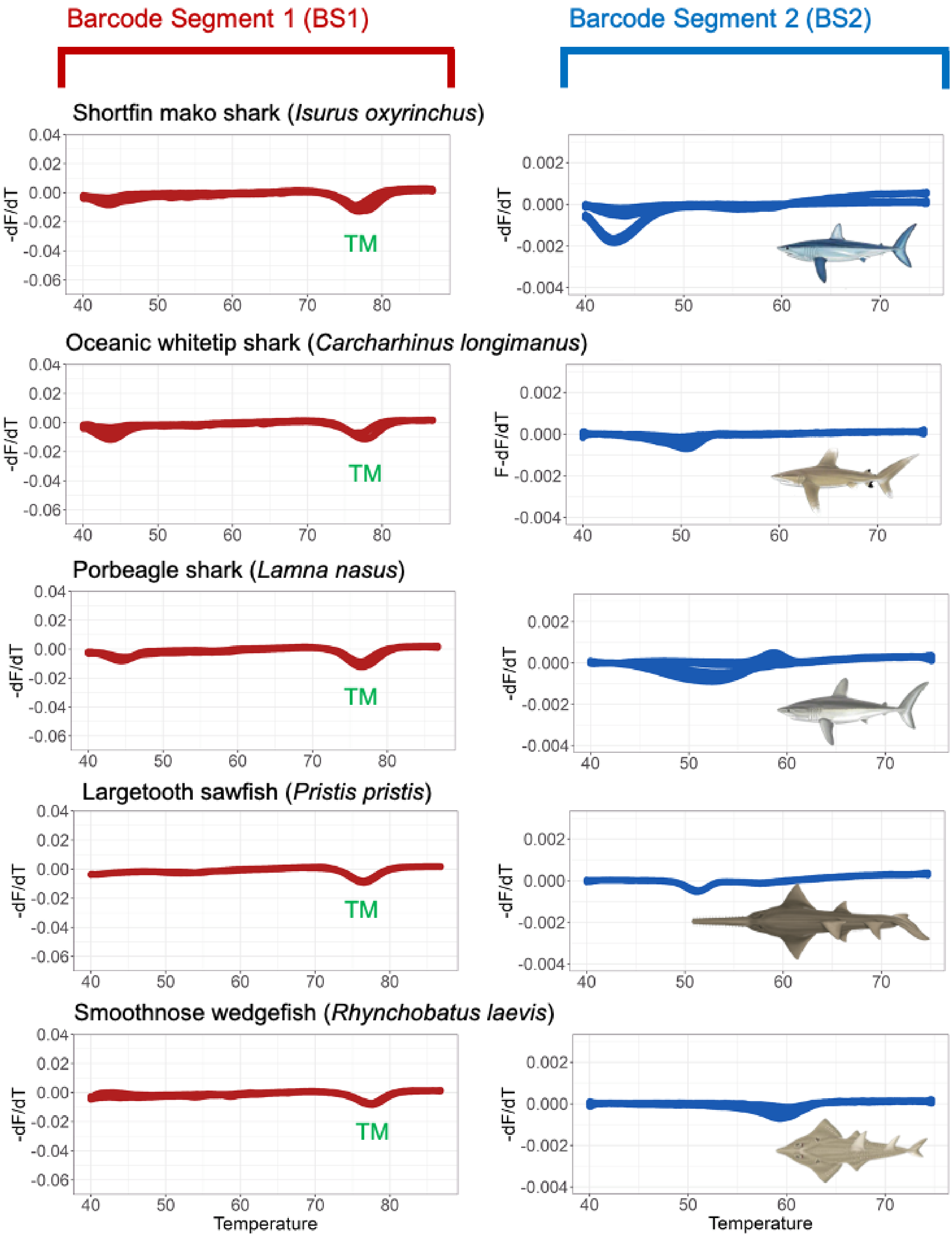


**Figure S.5**. Some species which have a hybridization problem in the BS1 region. Those species only have "TM" signature (the right-most valley in the BS1, labelled with a green color), TM corresponds to ThermaMark$^{TM}$, an internal marker for correction of artefactual temperature variation.

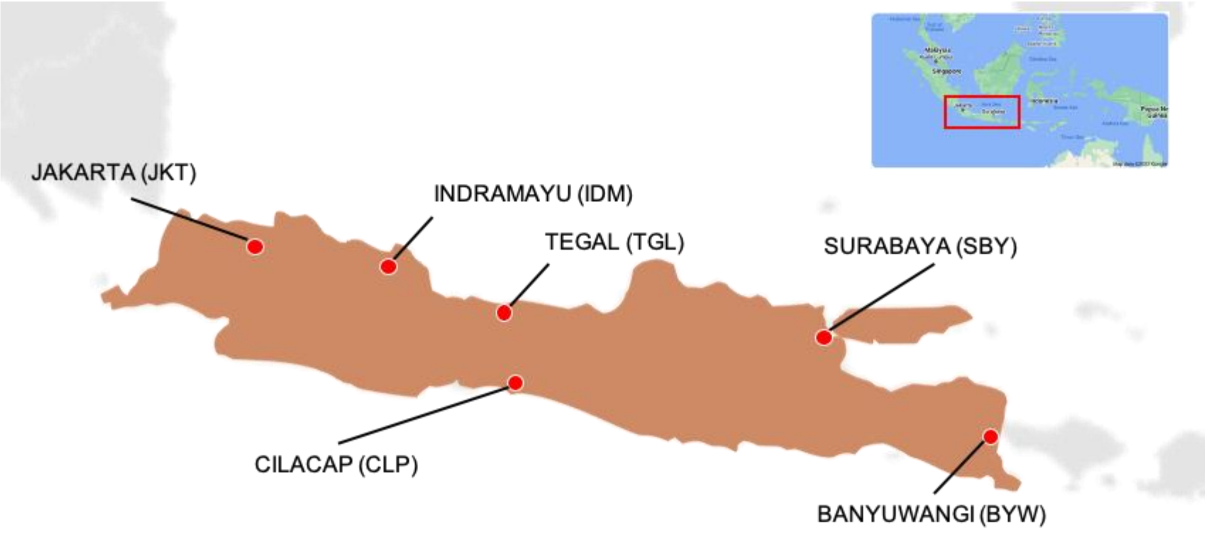


**Figure S.6**. Sampling locations across Java Island, Indonesia. Locations are labelled with long and short codes.

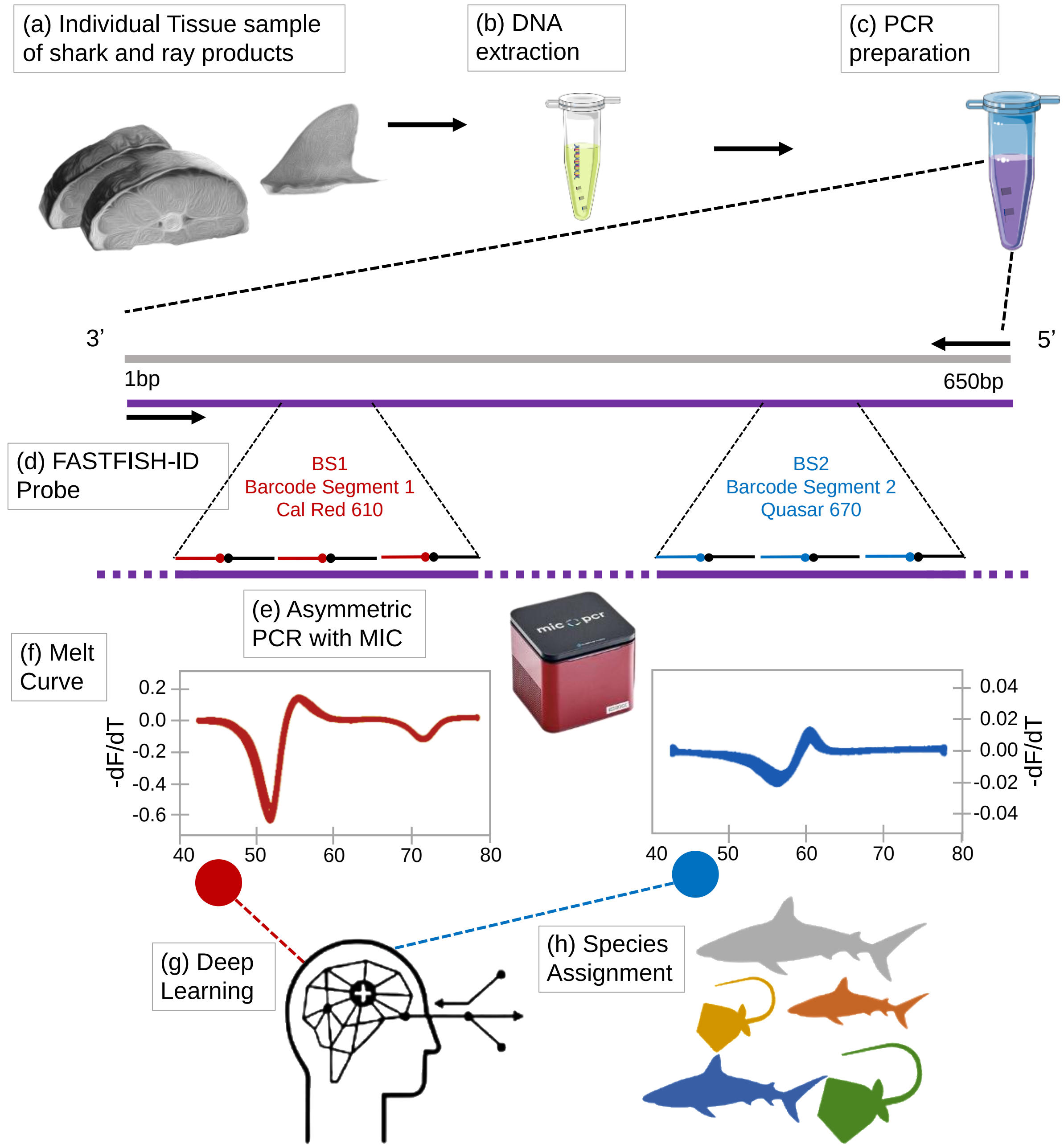


**Figure S.7**. A schematic description of the stages of this study which include (a) sample collection and preservation, (b) DNA extraction of tissue samples, (c-e) sample processing using the FASTFISH-ID workflow, (f) visualisation of the RT-PCR outputs and (g and h) species classification using deep learning.

### Supplementary tables

**Table S.1**. Sample details used on the training datasets including Condition (processed/fresh), Part (of the animal), Species, ID (number), no. of replications and Sequencing technology used to identify the species.

| Condition | Part | Species | ID | Replication | Sequencing |
|---|---|---|---|---|---|
| Processed | Dried fin | *Alopias pelagicus* | 340 | 3 | Sanger ~650bp |
| Processed | Dried fin | *Alopias pelagicus* | 341 | 2 | Sanger ~650bp |
| Processed | Dried fin | *Alopias superciliosus* | 54 | 3 | HTB ~313bp |
| Processed | Dried fin | *Alopias superciliosus* | 345 | 3 | Sanger ~650bp |
| Processed | Dried fin | *Alopias superciliosus* | 346 | 3 | Sanger ~650bp |
| Processed | Salted meat | *Alopias superciliosus* | 366 | 3 | HTB ~313bp |
| Processed | Dried fin | *Alopias superciliosus* | 431 | 2 | Sanger ~650bp |
| Processed | Unidentified | *Alopias superciliosus* | 530 | 3 | HTB ~313bp |
| Processed | Rostrum | *Anoxypristis cuspidata* | 9 | 4 | Sanger ~650bp |
| Processed | Dried fin | *Anoxypristis cuspidata* | 22 | 3 | Sanger ~650bp |
| Processed | Unidentified | *Anoxypristis cuspidata* | 536 | 3 | HTB ~313bp |
| Processed | Rostrum | *Anoxypristis cuspidata* | 490 | 2 | Sanger ~650bp |
| Fresh | Trunk | *Carcharhinus brevipinna* | 77 | 3 | HTB ~313bp |
| Fresh | Trunk | *Carcharhinus brevipinna* | 78 | 3 | Sanger ~650bp |
| Fresh | Trunk | *Carcharhinus brevipinna* | 86 | 2 | Sanger ~650bp |
| Fresh | Finless | *Carcharhinus brevipinna* | 123 | 3 | HTB ~313bp |
| Fresh | Whole | *Carcharhinus brevipinna* | 321 | 1 | Sanger ~650bp |
| Fresh | Whole | *Carcharhinus brevipinna* | 323 | 3 | Sanger ~650bp |
| Fresh | Whole | *Carcharhinus brevipinna* | 324 | 3 | Sanger ~650bp |
| Fresh | Whole | *Carcharhinus brevipinna* | 334 | 3 | Sanger ~650bp |
| **Condition** | **Part** | **Species** | **ID** | **Replication** | **Sequencing** |

| | | | | | |
|---|---|---|---|---|---|
| Fresh | Whole | *Carcharhinus brevipinna* | 475 | 1 | Sanger ~650bp |
| Fresh | Trunk | *Carcharhinus falciformis* | 3 | 3 | HTB ~313bp |
| Fresh | Trunk | *Carcharhinus falciformis* | 4 | 3 | HTB ~313bp |
| Fresh | Trunk | *Carcharhinus falciformis* | 5 | 3 | HTB ~313bp |
| Fresh | Trunk | *Carcharhinus falciformis* | 6 | 3 | HTB ~313bp |
| Fresh | Trunk | *Carcharhinus falciformis* | 7 | 3 | HTB ~313bp |
| Fresh | Trunk | *Carcharhinus falciformis* | 43 | 3 | Sanger ~650bp |
| Fresh | Whole | *Carcharhinus falciformis* | 285 | 3 | Sanger ~650bp |
| Fresh | Whole | *Carcharhinus falciformis* | 293 | 2 | Sanger ~650bp |
| Fresh | Whole | *Carcharhinus falciformis* | 294X | 3 | Sanger ~650bp |
| Processed | Dried fin | *Carcharhinus longimanus* | 25 | 3 | Sanger ~650bp |
| Processed | Dried fin | *Carcharhinus longimanus* | 53 | 3 | Sanger ~650bp |
| Processed | Dried fin | *Carcharhinus longimanus* | 342 | 2 | Sanger ~650bp |
| Fresh | Trunk | *Carcharhinus sorrah* | 29 | 3 | HTB ~313bp |
| Fresh | Trunk | *Carcharhinus sorrah* | 46 | 3 | HTB ~313bp |
| Fresh | Whole | *Carcharhinus sorrah* | 185 | 3 | Sanger ~650bp |
| Fresh | Whole | *Carcharhinus sorrah* | 319 | 1 | Sanger ~650bp |
| Fresh | Whole | *Galeocerdo cuvier* | 178 | 3 | HTB ~313bp |
| Fresh | Whole | *Galeocerdo cuvier* | 363 | 3 | HTB ~313bp |
| Fresh | Fin | *Galeocerdo cuvier* | 456 | 1 | Sanger ~650bp |
| Processed | Dried fin | *Galeocerdo cuvier* | 354 | 3 | Sanger ~650bp |
| Processed | Dried fin | *Galeocerdo cuvier* | 435 | 3 | Sanger ~650bp |
| Processed | Dried fin | *Galeocerdo cuvier* | 436 | 3 | Sanger ~650bp |
| Processed | Dried fin | *Galeocerdo cuvier* | 437 | 3 | Sanger ~650bp |
| **Condition** | **Part** | **Species** | **ID** | **Replication** | **Sequencing** |

| | | | | | |
|---|---|---|---|---|---|
| Processed | Teeth | *Galeocerdo cuvier* | 439 | 3 | Sanger ~650bp |
| Fresh | Whole | *Glaucostegus typus* | 212 | 3 | HTB ~313bp |
| Fresh | Whole | *Glaucostegus typus* | 268 | 5 | Sanger ~650bp |
| Processed | Dried fin | *Glaucostegus typus* | 11 | 3 | HTB ~313bp |
| Processed | Dried skin | *Glaucostegus typus* | 196 | 3 | HTB ~313bp |
| Fresh | Whole | *Gymnura poecilura* | 90 | 3 | Sanger ~650bp |
| Fresh | Whole | *Gymnura poecilura* | 91 | 3 | Sanger ~650bp |
| Fresh | Whole | *Gymnura poecilura* | 92 | 3 | Sanger ~650bp |
| Fresh | Whole | *Himantura imbricata* | 296 | 3 | Sanger ~650bp |
| Fresh | Whole | *Himantura imbricata* | 297 | 2 | Sanger ~650bp |
| Processed | Dried fin | *Isurus oxyrinchus* | 50 | 3 | Sanger ~650bp |
| Processed | Dried fin | *Isurus oxyrinchus* | 343 | 3 | Sanger ~650bp |
| Processed | Dried fin | *Isurus oxyrinchus* | 344 | 2 | Sanger ~650bp |
| Processed | Dried fin | *Isurus oxyrinchus* | 384 | 3 | HTB ~313bp |
| Processed | Dried fin | *Isurus oxyrinchus* | 421 | 3 | HTB ~313bp |
| Processed | Unidentified | *Isurus oxyrinchus* | 519 | 3 | Sanger ~650bp |
| Processed | Unidentified | *Isurus oxyrinchus* | 521 | 2 | Sanger ~650bp |
| Processed | Dried fin | *Isurus paucus* | 20 | 3 | Sanger ~650bp |
| Processed | Dried fin | *Isurus paucus* | 52 | 3 | Sanger ~650bp |
| Processed | Dried fin | *Isurus paucus* | 338 | 3 | Sanger ~650bp |
| Processed | Dried fin | *Isurus paucus* | 339 | 2 | Sanger ~650bp |
| Processed | Unidentified | *Isurus paucus* | 528 | 3 | HTB ~313bp |
| Processed | Unidentified | *Isurus paucus* | 533 | 3 | HTB ~313bp |
| Processed | Dried fin | *Lamna nasus* | 24 | 3 | HTB ~313bp |
| Processed | Dried fin | *Lamna nasus* | 505 | 3 | HTB ~313bp |
| Processed | Dried fin | *Lamna nasus* | 506 | 3 | HTB ~313bp |
| Processed | Unidentified | *Lamna nasus* | 527 | 3 | HTB ~313bp |
| Processed | Salted meat | *Mobula birostris* | 370 | 3 | HTB ~313bp |
| **Condition** | **Part** | **Species** | **ID** | **Replication** | **Sequencing** |

| | | | | | |
|---|---|---|---|---|---|
| Processed | Gill racker | *Mobula birostris* | 412 | 3 | HTB ~313bp |
| Processed | Gill racker | *Mobula mobular* | 448 | 3 | HTB ~313bp |
| Processed | Gill racker | *Mobula mobular* | 449 | 3 | HTB ~313bp |
| Processed | Gill racker | *Mobula mobular* | 450 | 3 | HTB ~313bp |
| Processed | Gill racker | *Mobula mobular* | 451 | 3 | HTB ~313bp |
| Processed | Cartillage | *Mobula tarapacana* | 12 | 3 | HTB ~313bp |
| Fresh | Whole | *Neotrygon orientalis* | 240 | 3 | Sanger ~650bp |
| Fresh | Whole | *Neotrygon orientalis* | 241 | 1 | Sanger ~650bp |
| Fresh | Whole | *Neotrygon orientalis* | 244 | 3 | Sanger ~650bp |
| Fresh | Trunk | *Prionace glauca* | 413 | 3 | Sanger ~650bp |
| Processed | Dried fin | *Prionace glauca* | 355 | 3 | Sanger ~650bp |
| Processed | Dried fin | *Prionace glauca* | 356 | 3 | Sanger ~650bp |
| Processed | Unidentified | *Pristis pristis* | 550 | 3 | HTB ~313bp |
| Fresh | Whole | *Rhina ancylostoma* | 276 | 3 | Sanger ~650bp |
| Fresh | Whole | *Rhina ancylostoma* | 211 | 3 | Sanger ~650bp |
| Processed | Dried fin | *Rhina ancylostoma* | 27 | 3 | Sanger ~650bp |
| Processed | Dried skin | *Rhina ancylostoma* | 48 | 4 | Sanger ~650bp |
| Processed | Meat | *Rhina ancylostoma* | 247 | 3 | HTB ~313bp |
| Fresh | Whole | *Rhynchobatus australiae* | 101 | 3 | Sanger ~650bp |
| Fresh | Finless | *Rhynchobatus australiae* | 175 | 3 | Sanger ~650bp |
| Fresh | Whole | *Rhynchobatus australiae* | 213 | 2 | Sanger ~650bp |
| Fresh | Whole | *Rhynchobatus australiae* | 229 | 1 | HTB ~313bp |
| Fresh | Whole | *Rhynchobatus australiae* | 259 | 1 | HTB ~313bp |
| Fresh | Whole | *Rhynchobatus australiae* | 279 | 3 | Sanger ~650bp |
| Processed | Dried fin | *Rhynchobatus australiae* | 424 | 3 | HTB ~313bp |
| Fresh | Whole | *Rhynchobatus laevis* | 35 | 3 | Sanger ~650bp |
| Fresh | Whole | *Rhynchobatus laevis* | 151 | 3 | Sanger ~650bp |
| Fresh | Whole | *Rhynchobatus laevis* | 152 | 3 | Sanger ~650bp |
| **Condition** | **Part** | **Species** | **ID** | **Replication** | **Sequencing** |

| | | | | | |
|---|---|---|---|---|---|
| Fresh | Finless | *Rhynchobatus laevis* | 177 | 3 | Sanger ~650bp |
| Fresh | Whole | *Rhynchobatus springeri* | 189 | 3 | Sanger ~650bp |
| Fresh | Whole | *Rhynchobatus springeri* | 214 | 3 | Sanger ~650bp |
| Fresh | Whole | *Rhynchobatus springeri* | 215 | 1 | HTB ~313bp |
| Fresh | Whole | *Rhynchobatus springeri* | 221 | 3 | Sanger ~650bp |
| Fresh | Whole | *Rhynchobatus springeri* | 224 | 1 | Sanger ~650bp |
| Fresh | Whole | *Rhynchobatus springeri* | 226 | 1 | Sanger ~650bp |
| Fresh | Whole | *Rhynchobatus springeri* | 258 | 3 | Sanger ~650bp |
| Fresh | Whole | *Rhynchobatus springeri* | 274 | 3 | Sanger ~650bp |
| Fresh | Finless | *Sphyrna lewini* | 112 | 3 | Sanger ~650bp |
| Fresh | Whole | *Sphyrna lewini* | 115 | 3 | Sanger ~650bp |
| Fresh | Whole | *Sphyrna lewini* | 121 | 3 | Sanger ~650bp |
| Fresh | Finless | *Sphyrna lewini* | 122 | 3 | HTB ~313bp |
| Fresh | Finless | *Sphyrna lewini* | 126 | 3 | Sanger ~650bp |
| Fresh | Whole | *Sphyrna lewini* | 476 | 3 | Sanger ~650bp |
| Processed | Dried fin | *Sphyrna lewini* | 16 | 3 | HTB ~313bp |
| Processed | Dried fin | *Sphyrna lewini* | 426 | 1 | Sanger ~650bp |
| Fresh | Finless | *Sphyrna mokarran* | 113 | 3 | Sanger ~650bp |
| Processed | Cartillage | *Sphyrna mokarran* | 13 | 3 | HTB ~313bp |
| Processed | Dried fin | *Sphyrna mokarran* | 21 | 3 | Sanger ~650bp |
| Processed | Dried skin | *Sphyrna mokarran* | 197 | 3 | HTB ~313bp |
| Processed | Salted meat | *Sphyrna mokarran* | 367 | 3 | HTB ~313bp |
| Processed | Dried fin | *Sphyrna mokarran* | 418 | 2 | Sanger ~650bp |
| Fresh | Whole | *Stegostoma fasciatum* | 133 | 3 | HTB ~313bp |
| Fresh | Trunk | *Stegostoma fasciatum* | 179 | 3 | HTB ~313bp |
| Fresh | Trunk | *Stegostoma fasciatum* | 180 | 3 | HTB ~313bp |

| Condition | Part | Species | ID | Replication | Sequencing |
|---|---|---|---|---|---|
| Fresh | Trunk | *Stegostoma fasciatum* | 181 | 3 | Sanger ~650bp |
| Processed | Dried fin | *Stegostoma fasciatum* | 583 | 1 | Sanger ~650bp |
| Fresh | Whole | *Telatrygon zugei* | 198 | 3 | Sanger ~650bp |
| Fresh | Whole | *Telatrygon zugei* | 245 | 2 | Sanger ~650bp |

**Table S.2**. Sample details used on the testing datasets including Condition (processed/fresh), Part (of the animal), Species, ID (number), no. of replications and Sequencing technology used to identify the species.

| Condition | Part | Species | ID | Replication | Sequencing |
|---|---|---|---|---|---|
| Processed | Dried fin | *Alopias pelagicus* | 340 | 1 | Sanger ~650bp |
| Processed | Dried fin | *Alopias superciliosus* | 431 | 1 | Sanger ~650bp |
| Processed | Unidentified | *Alopias superciliosus* | 535 | 1 | HTB ~313bp |
| Processed | Unidentified | *Anoxypristis cuspidata* | 536 | 1 | HTB ~313bp |
| Fresh | Whole | *Carcharhinus brevipinna* | 317 | 1 | HTB ~313bp |
| Fresh | Whole | *Carcharhinus brevipinna* | 321 | 1 | Sanger ~650bp |
| Fresh | Whole | *Carcharhinus brevipinna* | 322 | 1 | HTB ~313bp |
| Fresh | Whole | *Carcharhinus brevipinna* | 326 | 1 | HTB ~313bp |
| Fresh | Whole | *Carcharhinus brevipinna* | 475 | 1 | Sanger ~650bp |
| Fresh | Trunk | *Carcharhinus falciformis* | 43 | 1 | Sanger ~650bp |
| Fresh | Trunk | *Carcharhinus falciformis* | 4 | 1 | HTB ~313bp |
| Fresh | Trunk | *Carcharhinus falciformis* | 19 | 1 | HTB ~313bp |
| Fresh | Trunk | *Carcharhinus falciformis* | 58 | 1 | HTB ~313bp |
| Processed | Dried fin | *Carcharhinus longimanus* | 342 | 1 | Sanger ~650bp |
| Processed | Unidentified | *Carcharhinus longimanus* | 522 | 1 | HTB ~313bp |
| Processed | Unidentified | *Carcharhinus longimanus* | 523 | 1 | HTB ~313bp |
| Processed | Unidentified | *Carcharhinus longimanus* | 524 | 1 | HTB ~313bp |
| Fresh | Whole | *Carcharhinus sorrah* | 304 | 1 | Sanger ~650bp |
| Processed | Oil | *Galeocerdo cuvier* | 396 | 1 | HTB ~313bp |
| Processed | Dried fin | *Galeocerdo cuvier* | 432 | 1 | HTB ~313bp |
| Processed | Dried fin | *Galeocerdo cuvier* | 433 | 1 | HTB ~313bp |
| Processed | Dried fin | *Galeocerdo cuvier* | 434 | 1 | HTB ~313bp |

| Condition | Part | Species | ID | Replication | Sequencing |
|---|---|---|---|---|---|
| Processed | Dried skin | *Galeocerdo cuvier* | 441 | 1 | Sanger ~650bp |
| Fresh | Whole | *Glaucostegus typus* | 272 | 1 | Sanger ~650bp |
| Fresh | Whole | *Glaucostegus typus* | 275 | 1 | HTB ~313bp |
| Processed | Dried fin | *Glaucostegus typus* | 422 | 1 | HTB ~313bp |
| Processed | Dried fin | *Glaucostegus typus* | 428 | 1 | HTB ~313bp |
| Processed | Unidentified | *Glaucostegus typus* | 537 | 1 | HTB ~313bp |
| Fresh | Whole | *Gymnura poecilura* | 88 | 1 | Sanger ~650bp |
| Fresh | Whole | *Gymnura poecilura* | 89 | 1 | Sanger ~650bp |
| Fresh | Whole | *Himantura imbricata* | 297 | 1 | Sanger ~650bp |
| Processed | Dried fin | *Isurus oxyrinchus* | 344 | 1 | Sanger ~650bp |
| Processed | Unidentified | *Isurus oxyrinchus* | 531 | 1 | HTB ~313bp |
| Processed | Dried fin | *Isurus paucus* | 339 | 1 | Sanger ~650bp |
| Processed | Dried fin | *Lamna nasus* | 24 | 1 | HTB ~313bp |
| Processed | Unidentified | *Lamna nasus* | 529 | 1 | HTB ~313bp |
| Processed | Salted meat | *Mobula birostris* | 370 | 1 | HTB ~313bp |
| Processed | Gill racker | *Mobula mobular* | 451 | 1 | HTB ~313bp |
| Processed | Cartillage | *Mobula tarapacana* | 12 | 1 | HTB ~313bp |
| Fresh | Whole | *Neotrygon orientalis* | 242 | 1 | Sanger ~650bp |
| Fresh | Trunk | *Prionace glauca* | 414 | 1 | HTB ~313bp |
| Fresh | Trunk | *Prionace glauca* | 416 | 1 | Sanger ~650bp |
| Fresh | Trunk | *Prionace glauca* | 417 | 1 | HTB ~313bp |
| Processed | Dried fin unskin | *Prionace glauca* | 399 | 1 | HTB ~313bp |
| Processed | Dried fin | *Prionace glauca* | 410 | 1 | HTB ~313bp |
| Processed | Unidentified | *Pristis pristis* | 550 | 1 | HTB ~313bp |
| Processed | Dried fin | *Rhina ancylostoma* | 14 | 1 | Sanger ~650bp |
| Processed | Dried skin | *Rhina ancylostoma* | 48 | 1 | Sanger ~650bp |
| Fresh | Whole | *Rhynchobatus australiae* | 213 | 1 | Sanger ~650bp |
| Fresh | Whole | *Rhynchobatus laevis* | 39 | 1 | Sanger ~650bp |
| Fresh | Finless | *Rhynchobatus laevis* | 176 | 1 | HTB ~313bp |

| Condition | Part | Species | ID | Replication | Sequencing |
|---|---|---|---|---|---|
| Processed | Unidentified | *Rhynchobatus laevis* | 534 | 1 | HTB ~313bp |
| Fresh | Whole | *Rhynchobatus springeri* | 217 | 1 | HTB ~313bp |
| Fresh | Whole | *Rhynchobatus springeri* | 224 | 1 | Sanger ~650bp |
| Fresh | Whole | *Rhynchobatus springeri* | 225 | 1 | Sanger ~650bp |
| Fresh | Whole | *Rhynchobatus springeri* | 226 | 1 | Sanger ~650bp |
| Fresh | Whole | *Rhynchobatus springeri* | 223B | 1 | Sanger ~650bp |
| Fresh | Finless | *Sphyrna lewini* | 125 | 1 | HTB ~313bp |
| Fresh | Trunk | *Sphyrna lewini* | 155 | 1 | HTB ~313bp |
| Fresh | Trunk | *Sphyrna lewini* | 156 | 1 | HTB ~313bp |
| Fresh | Whole | *Sphyrna lewini* | 160 | 1 | HTB ~313bp |
| Fresh | Whole | *Sphyrna lewini* | 234 | 1 | Sanger ~650bp |
| Processed | Dried fin | *Sphyrna lewini* | 419 | 1 | Sanger ~650bp |
| Processed | Dried fin | *Sphyrna lewini* | 426 | 1 | Sanger ~650bp |
| Processed | Dried fin | *Sphyrna mokarran* | 418 | 1 | Sanger ~650bp |
| Processed | Dried fin | *Sphyrna mokarran* | 420 | 1 | HTB ~313bp |
| Processed | Dried skin | *Stegostoma fasciatum* | 195 | 1 | HTB ~313bp |
| Fresh | Whole | *Telatrygon zugei* | 245 | 1 | Sanger ~650bp |

**Table S.3**. Variable importance in recognizing fluorescent signatures of species

| Barcode segment | Variable | Relative importance | Scaled importance | Percentage |
|---|---|---|---|---|
| BS1 | C5 | 1 | 1 | 1.87E-04 |
| BS1 | C13 | 0.97 | 0.97 | 1.81E-04 |
| BS1 | C15 | 0.96 | 0.96 | 1.80E-04 |
| BS1 | C17 | 0.97 | 0.97 | 1.82E-04 |
| … | … | … | … | … |
| BS1 | C2635 | 0.53 | 0.53 | 9.90E-05 |
| BS2 | C4678 | 0.98 | 0.98 | 1.82E-04 |
| BS2 | C6741 | 0.52 | 0.52 | 9.81E-05 |
| BS2 | C6747 | 0.53 | 0.53 | 9.92E-05 |
| BS2 | C6748 | 0.53 | 0.53 | 9.91E-05 |
| BS2 | C6750 | 0.53 | 0.53 | 9.90E-05 |

**Table S.4**. Result of grid search in finding the best deep learning model

| No | Model ID | Accuracy | Activation function | Epochs | Epsilon | Hidden layers | Input dropout ratio | L1 | L2 | Max w2 | Rho |
|---|---|---|---|---|---|---|---|---|---|---|---|
| 1 | dl_grid_model_17 | 0.98 | RectifierWithDropout | 500 | 1.00E-08 | [500, 500, 500] | 0.2 | 0 | 0.0001 | 1000 | 0.9 |
| 2 | dl_grid_model_170 | 0.98 | Maxout | 300 | 1.00E-06 | [500, 500, 500] | 0.2 | 0 | 0 | 100 | 0.9 |
| 3 | dl_grid_model_7 | 0.98 | MaxoutWithDropout | 500 | 1.00E-06 | [100, 100, 100] | 0 | 0 | 0.0001 | 100 | 0.95 |
| 4 | dl_grid_model_104 | 0.97 | Tanh | 500 | 1.00E-10 | [100, 100, 100] | 0.2 | 0 | 0 | 10 | 0.95 |
| 5 | dl_grid_model_107 | 0.97 | TanhWithDropout | 300 | 1.00E-06 | [500, 500, 500] | 0 | 1E-05 | 1E-05 | 1000 | 0.95 |
| ... | ... | ... | ... | ... | ... | ... | ... | ... | ... | ... | ... |
| 7 | dl_grid_model_32 | 0.01 | Rectifier | 300 | 1.00E-04 | [100, 100, 100] | 0.1 | 0 | 0 | 10 | 1 |
| 8 | dl_grid_model_195 | 0.00 | Rectifier | 100 | 1.00E-04 | [500, 500, 500] | 0 | 0 | 0 | 10 | 1 |
| 9 | dl_grid_model_247 | 0.00 | RectifierWithDropout | 200 | 1.00E-06 | [500, 500, 500] | 0 | 0 | 0 | 1000 | 1 |
| 10 | dl_grid_model_260 | 0.00 | RectifierWithDropout | 300 | 1.00E-04 | [200, 200, 200] | 0 | 0 | 1E-05 | 100 | 0.95 |
| 11 | dl_grid_model_66 | 0.00 | RectifierWithDropout | 50 | 1.00E-04 | [200, 200, 200] | 0 | 0 | 0.0001 | 100 | 0.95 |

**Table S.5**. Assignment scoring of 28 species of sharks and rays

| No. | Actual | Prediction | | SCORE | *Alopias_pelagicus* | *Alopias_superciliosus* | *Anoxypristis_cuspidata* | *Carcharhinus_brevipinna* | *Carcharhinus_falciformis* | *Carcharhinus_longimanus* | *Carcharhinus_sorrah* | *Galeocerdo_cuvier* | *Glaucostegus_typus* | *Gymnura_poecilura* | *Himantura_imbricata* | *Isurus_oxyrinchus* | *Isurus_paucus* | *Lamna_nasus* | *Mobula_birostris* | *Mobula_mobular* | *Mobula_tarapacana* | *Neotrygon_orientalis* | *Prionace_glauca* | *Pristis_pristis* | *Rhina_ancylostoma* | *Rhynchobatus_australiae* | *Rhynchobatus_laevis* | *Rhynchobatus_springeri* | *Sphyrna_lewini* | *Sphyrna_mokarran* | *Stegostoma_fasciatum* | *Telatrygon_zugei* |
|---|---|---|---|---|---|---|---|---|---|---|---|---|---|---|---|---|---|---|---|---|---|---|---|---|---|---|---|---|---|---|---|---|
| 1 | *Glaucostegus_typus* | *Glaucostegus_typus* | Match | 1.000 | 0.000 | 0.000 | 0.000 | 0.000 | 0.000 | 0.000 | 0.000 | 0.000 | 1.000 | 0.000 | 0.000 | 0.000 | 0.000 | 0.000 | 0.000 | 0.000 | 0.000 | 0.000 | 0.000 | 0.000 | 0.000 | 0.000 | 0.000 | 0.000 | 0.000 | 0.000 | 0.000 | 0.000 |
| 2 | *Rhina_ancylostoma* | *Rhina_ancylostoma* | Match | 1.000 | 0.000 | 0.000 | 0.000 | 0.000 | 0.000 | 0.000 | 0.000 | 0.000 | 0.000 | 0.000 | 0.000 | 0.000 | 0.000 | 0.000 | 0.000 | 0.000 | 0.000 | 0.000 | 0.000 | 0.000 | 1.000 | 0.000 | 0.000 | 0.000 | 0.000 | 0.000 | 0.000 | 0.000 |
| 3 | *Rhynchobatus_laevis* | *Rhynchobatus_laevis* | Match | 0.545 | 0.004 | 0.001 | 0.000 | 0.000 | 0.001 | 0.000 | 0.001 | 0.319 | 0.102 | 0.000 | 0.000 | 0.000 | 0.000 | 0.000 | 0.000 | 0.000 | 0.001 | 0.000 | 0.002 | 0.000 | 0.002 | 0.001 | 0.545 | 0.000 | 0.019 | 0.000 | 0.000 | 0.000 |
| 4 | *Rhynchobatus_springeri* | *Rhynchobatus_springeri* | Match | 0.999 | 0.000 | 0.000 | 0.000 | 0.000 | 0.000 | 0.000 | 0.000 | 0.000 | 0.000 | 0.000 | 0.000 | 0.000 | 0.000 | 0.000 | 0.000 | 0.000 | 0.000 | 0.000 | 0.000 | 0.000 | 0.000 | 0.000 | 0.001 | 0.999 | 0.000 | 0.000 | 0.000 | 0.000 |
| 5 | *Rhynchobatus_springeri* | *Rhynchobatus_springeri* | Match | 1.000 | 0.000 | 0.000 | 0.000 | 0.000 | 0.000 | 0.000 | 0.000 | 0.000 | 0.000 | 0.000 | 0.000 | 0.000 | 0.000 | 0.000 | 0.000 | 0.000 | 0.000 | 0.000 | 0.000 | 0.000 | 0.000 | 0.000 | 0.000 | 1.000 | 0.000 | 0.000 | 0.000 | 0.000 |
| 6 | *Gymnura_poecilura* | *Gymnura_poecilura* | Match | 1.000 | 0.000 | 0.000 | 0.000 | 0.000 | 0.000 | 0.000 | 0.000 | 0.000 | 0.000 | 1.000 | 0.000 | 0.000 | 0.000 | 0.000 | 0.000 | 0.000 | 0.000 | 0.000 | 0.000 | 0.000 | 0.000 | 0.000 | 0.000 | 0.000 | 0.000 | 0.000 | 0.000 | 0.000 |
| 7 | *Gymnura_poecilura* | *Gymnura_poecilura* | Match | 1.000 | 0.000 | 0.000 | 0.000 | 0.000 | 0.000 | 0.000 | 0.000 | 0.000 | 0.000 | 1.000 | 0.000 | 0.000 | 0.000 | 0.000 | 0.000 | 0.000 | 0.000 | 0.000 | 0.000 | 0.000 | 0.000 | 0.000 | 0.000 | 0.000 | 0.000 | 0.000 | 0.000 | 0.000 |
| 8 | *Neotrygon_orientalis* | *Neotrygon_orientalis* | Match | 1.000 | 0.000 | 0.000 | 0.000 | 0.000 | 0.000 | 0.000 | 0.000 | 0.000 | 0.000 | 0.000 | 0.000 | 0.000 | 0.000 | 0.000 | 0.000 | 0.000 | 0.000 | 1.000 | 0.000 | 0.000 | 0.000 | 0.000 | 0.000 | 0.000 | 0.000 | 0.000 | 0.000 | 0.000 |
| 9 | *Sphyrna_lewini* | *Sphyrna_lewini* | Match | 1.000 | 0.000 | 0.000 | 0.000 | 0.000 | 0.000 | 0.000 | 0.000 | 0.000 | 0.000 | 0.000 | 0.000 | 0.000 | 0.000 | 0.000 | 0.000 | 0.000 | 0.000 | 0.000 | 0.000 | 0.000 | 0.000 | 0.000 | 0.000 | 0.000 | 1.000 | 0.000 | 0.000 | 0.000 |
| 10 | *Sphyrna_lewini* | *Sphyrna_lewini* | Match | 1.000 | 0.000 | 0.000 | 0.000 | 0.000 | 0.000 | 0.000 | 0.000 | 0.000 | 0.000 | 0.000 | 0.000 | 0.000 | 0.000 | 0.000 | 0.000 | 0.000 | 0.000 | 0.000 | 0.000 | 0.000 | 0.000 | 0.000 | 0.000 | 0.000 | 1.000 | 0.000 | 0.000 | 0.000 |
| 11 | *Carcharhinus_sorrah* | *Stegostoma_fasciatum* | Mismatch | 0.975 | 0.000 | 0.000 | 0.000 | 0.000 | 0.000 | 0.000 | 0.025 | 0.000 | 0.000 | 0.000 | 0.000 | 0.000 | 0.000 | 0.000 | 0.000 | 0.000 | 0.000 | 0.000 | 0.000 | 0.000 | 0.000 | 0.000 | 0.000 | 0.000 | 0.000 | 0.000 | 0.975 | 0.000 |
| 12 | *Galeocerdo_cuvier* | *Galeocerdo_cuvier* | Match | 1.000 | 0.000 | 0.000 | 0.000 | 0.000 | 0.000 | 0.000 | 0.000 | 1.000 | 0.000 | 0.000 | 0.000 | 0.000 | 0.000 | 0.000 | 0.000 | 0.000 | 0.000 | 0.000 | 0.000 | 0.000 | 0.000 | 0.000 | 0.000 | 0.000 | 0.000 | 0.000 | 0.000 | 0.000 |
| 13 | *Prionace_glauca* | *Prionace_glauca* | Match | 1.000 | 0.000 | 0.000 | 0.000 | 0.000 | 0.000 | 0.000 | 0.000 | 0.000 | 0.000 | 0.000 | 0.000 | 0.000 | 0.000 | 0.000 | 0.000 | 0.000 | 0.000 | 0.000 | 1.000 | 0.000 | 0.000 | 0.000 | 0.000 | 0.000 | 0.000 | 0.000 | 0.000 | 0.000 |
| 14 | *Rhynchobatus_springeri* | *Rhynchobatus_springeri* | Match | 1.000 | 0.000 | 0.000 | 0.000 | 0.000 | 0.000 | 0.000 | 0.000 | 0.000 | 0.000 | 0.000 | 0.000 | 0.000 | 0.000 | 0.000 | 0.000 | 0.000 | 0.000 | 0.000 | 0.000 | 0.000 | 0.000 | 0.000 | 0.000 | 1.000 | 0.000 | 0.000 | 0.000 | 0.000 |
| 15 | *Rhynchobatus_springeri* | *Rhynchobatus_springeri* | Match | 1.000 | 0.000 | 0.000 | 0.000 | 0.000 | 0.000 | 0.000 | 0.000 | 0.000 | 0.000 | 0.000 | 0.000 | 0.000 | 0.000 | 0.000 | 0.000 | 0.000 | 0.000 | 0.000 | 0.000 | 0.000 | 0.000 | 0.000 | 0.000 | 1.000 | 0.000 | 0.000 | 0.000 | 0.000 |
| 16 | *Sphyrna_lewini* | *Sphyrna_lewini* | Match | 1.000 | 0.000 | 0.000 | 0.000 | 0.000 | 0.000 | 0.000 | 0.000 | 0.000 | 0.000 | 0.000 | 0.000 | 0.000 | 0.000 | 0.000 | 0.000 | 0.000 | 0.000 | 0.000 | 0.000 | 0.000 | 0.000 | 0.000 | 0.000 | 0.000 | 1.000 | 0.000 | 0.000 | 0.000 |
| 17 | *Carcharhinus_brevipinna* | *Carcharhinus_brevipinna* | Match | 1.000 | 0.000 | 0.000 | 0.000 | 1.000 | 0.000 | 0.000 | 0.000 | 0.000 | 0.000 | 0.000 | 0.000 | 0.000 | 0.000 | 0.000 | 0.000 | 0.000 | 0.000 | 0.000 | 0.000 | 0.000 | 0.000 | 0.000 | 0.000 | 0.000 | 0.000 | 0.000 | 0.000 | 0.000 |
| 18 | *Carcharhinus_brevipinna* | *Carcharhinus_brevipinna* | Match | 1.000 | 0.000 | 0.000 | 0.000 | 1.000 | 0.000 | 0.000 | 0.000 | 0.000 | 0.000 | 0.000 | 0.000 | 0.000 | 0.000 | 0.000 | 0.000 | 0.000 | 0.000 | 0.000 | 0.000 | 0.000 | 0.000 | 0.000 | 0.000 | 0.000 | 0.000 | 0.000 | 0.000 | 0.000 |
| 19 | *Rhina_ancylostoma* | *Rhina_ancylostoma* | Match | 1.000 | 0.000 | 0.000 | 0.000 | 0.000 | 0.000 | 0.000 | 0.000 | 0.000 | 0.000 | 0.000 | 0.000 | 0.000 | 0.000 | 0.000 | 0.000 | 0.000 | 0.000 | 0.000 | 0.000 | 0.000 | 1.000 | 0.000 | 0.000 | 0.000 | 0.000 | 0.000 | 0.000 | 0.000 |
| 20 | *Alopias_superciliosus* | *Alopias_superciliosus* | Match | 1.000 | 0.000 | 1.000 | 0.000 | 0.000 | 0.000 | 0.000 | 0.000 | 0.000 | 0.000 | 0.000 | 0.000 | 0.000 | 0.000 | 0.000 | 0.000 | 0.000 | 0.000 | 0.000 | 0.000 | 0.000 | 0.000 | 0.000 | 0.000 | 0.000 | 0.000 | 0.000 | 0.000 | 0.000 |
| 21 | *Carcharhinus_longimanus* | *Carcharhinus_longimanus* | Match | 1.000 | 0.000 | 0.000 | 0.000 | 0.000 | 0.000 | 1.000 | 0.000 | 0.000 | 0.000 | 0.000 | 0.000 | 0.000 | 0.000 | 0.000 | 0.000 | 0.000 | 0.000 | 0.000 | 0.000 | 0.000 | 0.000 | 0.000 | 0.000 | 0.000 | 0.000 | 0.000 | 0.000 | 0.000 |
| 22 | *Himantura_imbricata* | *Himantura_imbricata* | Match | 1.000 | 0.000 | 0.000 | 0.000 | 0.000 | 0.000 | 0.000 | 0.000 | 0.000 | 0.000 | 0.000 | 1.000 | 0.000 | 0.000 | 0.000 | 0.000 | 0.000 | 0.000 | 0.000 | 0.000 | 0.000 | 0.000 | 0.000 | 0.000 | 0.000 | 0.000 | 0.000 | 0.000 | 0.000 |
| 23 | *Isurus_oxyrinchus* | *Isurus_oxyrinchus* | Match | 1.000 | 0.000 | 0.000 | 0.000 | 0.000 | 0.000 | 0.000 | 0.000 | 0.000 | 0.000 | 0.000 | 0.000 | 1.000 | 0.000 | 0.000 | 0.000 | 0.000 | 0.000 | 0.000 | 0.000 | 0.000 | 0.000 | 0.000 | 0.000 | 0.000 | 0.000 | 0.000 | 0.000 | 0.000 |
| 24 | *Isurus_paucus* | *Isurus_paucus* | Match | 0.997 | 0.000 | 0.000 | 0.000 | 0.000 | 0.000 | 0.000 | 0.002 | 0.000 | 0.000 | 0.000 | 0.000 | 0.000 | 0.997 | 0.000 | 0.000 | 0.000 | 0.000 | 0.000 | 0.000 | 0.000 | 0.000 | 0.000 | 0.000 | 0.000 | 0.000 | 0.000 | 0.000 | 0.000 |
| 25 | *Rhynchobatus_australiae* | *Rhynchobatus_australiae* | Match | 1.000 | 0.000 | 0.000 | 0.000 | 0.000 | 0.000 | 0.000 | 0.000 | 0.000 | 0.000 | 0.000 | 0.000 | 0.000 | 0.000 | 0.000 | 0.000 | 0.000 | 0.000 | 0.000 | 0.000 | 0.000 | 0.000 | 1.000 | 0.000 | 0.000 | 0.000 | 0.000 | 0.000 | 0.000 |
| 26 | *Sphyrna_mokarran* | *Glaucostegus_typus* | Mismatch | 0.802 | 0.002 | 0.000 | 0.000 | 0.001 | 0.000 | 0.190 | 0.000 | 0.000 | 0.802 | 0.000 | 0.001 | 0.000 | 0.000 | 0.000 | 0.000 | 0.000 | 0.000 | 0.000 | 0.000 | 0.001 | 0.001 | 0.001 | 0.000 | 0.000 | 0.000 | 0.000 | 0.000 | 0.000 |
| 27 | *Telatrygon_zugei* | *Telatrygon_zugei* | Match | 1.000 | 0.000 | 0.000 | 0.000 | 0.000 | 0.000 | 0.000 | 0.000 | 0.000 | 0.000 | 0.000 | 0.000 | 0.000 | 0.000 | 0.000 | 0.000 | 0.000 | 0.000 | 0.000 | 0.000 | 0.000 | 0.000 | 0.000 | 0.000 | 0.000 | 0.000 | 0.000 | 0.000 | 1.000 |
| 28 | *Carcharhinus_brevipinna* | *Carcharhinus_brevipinna* | Match | 1.000 | 0.000 | 0.000 | 0.000 | 1.000 | 0.000 | 0.000 | 0.000 | 0.000 | 0.000 | 0.000 | 0.000 | 0.000 | 0.000 | 0.000 | 0.000 | 0.000 | 0.000 | 0.000 | 0.000 | 0.000 | 0.000 | 0.000 | 0.000 | 0.000 | 0.000 | 0.000 | 0.000 | 0.000 |
| 29 | *Carcharhinus_brevipinna* | *Carcharhinus_brevipinna* | Match | 1.000 | 0.000 | 0.000 | 0.000 | 1.000 | 0.000 | 0.000 | 0.000 | 0.000 | 0.000 | 0.000 | 0.000 | 0.000 | 0.000 | 0.000 | 0.000 | 0.000 | 0.000 | 0.000 | 0.000 | 0.000 | 0.000 | 0.000 | 0.000 | 0.000 | 0.000 | 0.000 | 0.000 | 0.000 |
| 30 | *Carcharhinus_brevipinna* | *Carcharhinus_brevipinna* | Match | 1.000 | 0.000 | 0.000 | 0.000 | 1.000 | 0.000 | 0.000 | 0.000 | 0.000 | 0.000 | 0.000 | 0.000 | 0.000 | 0.000 | 0.000 | 0.000 | 0.000 | 0.000 | 0.000 | 0.000 | 0.000 | 0.000 | 0.000 | 0.000 | 0.000 | 0.000 | 0.000 | 0.000 | 0.000 |
| 31 | *Carcharhinus_falciformis* | *Carcharhinus_falciformis* | Match | 0.995 | 0.000 | 0.000 | 0.000 | 0.005 | 0.995 | 0.000 | 0.000 | 0.000 | 0.000 | 0.000 | 0.000 | 0.000 | 0.000 | 0.000 | 0.000 | 0.000 | 0.000 | 0.000 | 0.000 | 0.000 | 0.000 | 0.000 | 0.000 | 0.000 | 0.000 | 0.000 | 0.000 | 0.000 |
| 32 | *Galeocerdo_cuvier* | *Glaucostegus_typus* | Mismatch | 1.000 | 0.000 | 0.000 | 0.000 | 0.000 | 0.000 | 0.000 | 0.000 | 0.000 | 1.000 | 0.000 | 0.000 | 0.000 | 0.000 | 0.000 | 0.000 | 0.000 | 0.000 | 0.000 | 0.000 | 0.000 | 0.000 | 0.000 | 0.000 | 0.000 | 0.000 | 0.000 | 0.000 | 0.000 |
| 33 | *Galeocerdo_cuvier* | *Glaucostegus_typus* | Mismatch | 0.923 | 0.000 | 0.000 | 0.000 | 0.000 | 0.000 | 0.000 | 0.000 | 0.050 | 0.923 | 0.000 | 0.000 | 0.026 | 0.000 | 0.000 | 0.000 | 0.000 | 0.000 | 0.000 | 0.000 | 0.000 | 0.000 | 0.000 | 0.000 | 0.000 | 0.000 | 0.000 | 0.000 | 0.000 |
| 34 | *Galeocerdo_cuvier* | *Glaucostegus_typus* | Mismatch | 1.000 | 0.000 | 0.000 | 0.000 | 0.000 | 0.000 | 0.000 | 0.000 | 0.000 | 1.000 | 0.000 | 0.000 | 0.000 | 0.000 | 0.000 | 0.000 | 0.000 | 0.000 | 0.000 | 0.000 | 0.000 | 0.000 | 0.000 | 0.000 | 0.000 | 0.000 | 0.000 | 0.000 | 0.000 |
| 35 | *Galeocerdo_cuvier* | *Glaucostegus_typus* | Mismatch | 1.000 | 0.000 | 0.000 | 0.000 | 0.000 | 0.000 | 0.000 | 0.000 | 0.000 | 1.000 | 0.000 | 0.000 | 0.000 | 0.000 | 0.000 | 0.000 | 0.000 | 0.000 | 0.000 | 0.000 | 0.000 | 0.000 | 0.000 | 0.000 | 0.000 | 0.000 | 0.000 | 0.000 | 0.000 |
| 36 | *Glaucostegus_typus* | *Glaucostegus_typus* | Match | 1.000 | 0.000 | 0.000 | 0.000 | 0.000 | 0.000 | 0.000 | 0.000 | 0.000 | 1.000 | 0.000 | 0.000 | 0.000 | 0.000 | 0.000 | 0.000 | 0.000 | 0.000 | 0.000 | 0.000 | 0.000 | 0.000 | 0.000 | 0.000 | 0.000 | 0.000 | 0.000 | 0.000 | 0.000 |
| 37 | *Glaucostegus_typus* | *Glaucostegus_typus* | Match | 1.000 | 0.000 | 0.000 | 0.000 | 0.000 | 0.000 | 0.000 | 0.000 | 0.000 | 1.000 | 0.000 | 0.000 | 0.000 | 0.000 | 0.000 | 0.000 | 0.000 | 0.000 | 0.000 | 0.000 | 0.000 | 0.000 | 0.000 | 0.000 | 0.000 | 0.000 | 0.000 | 0.000 | 0.000 |
| 38 | *Glaucostegus_typus* | *Glaucostegus_typus* | Match | 1.000 | 0.000 | 0.000 | 0.000 | 0.000 | 0.000 | 0.000 | 0.000 | 0.000 | 1.000 | 0.000 | 0.000 | 0.000 | 0.000 | 0.000 | 0.000 | 0.000 | 0.000 | 0.000 | 0.000 | 0.000 | 0.000 | 0.000 | 0.000 | 0.000 | 0.000 | 0.000 | 0.000 | 0.000 |
| 39 | *Glaucostegus_typus* | *Glaucostegus_typus* | Match | 1.000 | 0.000 | 0.000 | 0.000 | 0.000 | 0.000 | 0.000 | 0.000 | 0.000 | 1.000 | 0.000 | 0.000 | 0.000 | 0.000 | 0.000 | 0.000 | 0.000 | 0.000 | 0.000 | 0.000 | 0.000 | 0.000 | 0.000 | 0.000 | 0.000 | 0.000 | 0.000 | 0.000 | 0.000 |
| 40 | *Prionace_glauca* | *Prionace_glauca* | Match | 0.929 | 0.000 | 0.000 | 0.000 | 0.071 | 0.000 | 0.000 | 0.000 | 0.000 | 0.000 | 0.000 | 0.000 | 0.000 | 0.000 | 0.000 | 0.000 | 0.000 | 0.000 | 0.000 | 0.929 | 0.000 | 0.000 | 0.000 | 0.000 | 0.000 | 0.000 | 0.000 | 0.000 | 0.000 |
| 41 | *Prionace_glauca* | *Carcharhinus_brevipinna* | Mismatch | 0.878 | 0.000 | 0.000 | 0.000 | 0.878 | 0.000 | 0.000 | 0.000 | 0.000 | 0.000 | 0.000 | 0.000 | 0.000 | 0.000 | 0.000 | 0.000 | 0.000 | 0.000 | 0.000 | 0.122 | 0.000 | 0.000 | 0.000 | 0.000 | 0.000 | 0.000 | 0.000 | 0.000 | 0.000 |
| 42 | *Prionace_glauca* | *Prionace_glauca* | Match | 0.663 | 0.000 | 0.000 | 0.000 | 0.337 | 0.000 | 0.000 | 0.000 | 0.000 | 0.000 | 0.000 | 0.000 | 0.000 | 0.000 | 0.000 | 0.000 | 0.000 | 0.000 | 0.000 | 0.663 | 0.000 | 0.000 | 0.000 | 0.000 | 0.000 | 0.000 | 0.000 | 0.000 | 0.000 |
| 43 | *Prionace_glauca* | *Prionace_glauca* | Match | 0.735 | 0.000 | 0.000 | 0.000 | 0.192 | 0.073 | 0.000 | 0.000 | 0.000 | 0.000 | 0.000 | 0.000 | 0.000 | 0.000 | 0.000 | 0.000 | 0.000 | 0.000 | 0.000 | 0.735 | 0.000 | 0.000 | 0.000 | 0.000 | 0.000 | 0.000 | 0.000 | 0.000 | 0.000 |
| 44 | *Rhynchobatus_laevis* | *Isurus_paucus* | Mismatch | 0.740 | 0.003 | 0.001 | 0.000 | 0.000 | 0.000 | 0.000 | 0.000 | 0.000 | 0.006 | 0.000 | 0.000 | 0.000 | 0.740 | 0.000 | 0.003 | 0.004 | 0.001 | 0.000 | 0.000 | 0.004 | 0.014 | 0.077 | 0.136 | 0.009 | 0.000 | 0.000 | 0.001 | 0.000 |
| 45 | *Rhynchobatus_laevis* | *Rhynchobatus_laevis* | Match | 1.000 | 0.000 | 0.000 | 0.000 | 0.000 | 0.000 | 0.000 | 0.000 | 0.000 | 0.000 | 0.000 | 0.000 | 0.000 | 0.000 | 0.000 | 0.000 | 0.000 | 0.000 | 0.000 | 0.000 | 0.000 | 0.000 | 0.000 | 1.000 | 0.000 | 0.000 | 0.000 | 0.000 | 0.000 |
| 46 | *Rhynchobatus_springeri* | *Pristis_pristis* | Mismatch | 0.583 | 0.000 | 0.369 | 0.002 | 0.000 | 0.000 | 0.000 | 0.000 | 0.000 | 0.002 | 0.001 | 0.003 | 0.000 | 0.001 | 0.000 | 0.003 | 0.024 | 0.005 | 0.000 | 0.002 | 0.583 | 0.003 | 0.000 | 0.001 | 0.000 | 0.000 | 0.000 | 0.000 | 0.000 |
| 47 | *Sphyrna_lewini* | *Mobula_tarapacana* | Mismatch | 0.724 | 0.000 | 0.000 | 0.000 | 0.000 | 0.000 | 0.000 | 0.000 | 0.000 | 0.000 | 0.000 | 0.000 | 0.000 | 0.000 | 0.000 | 0.000 | 0.000 | 0.724 | 0.000 | 0.000 | 0.000 | 0.000 | 0.000 | 0.000 | 0.000 | 0.276 | 0.000 | 0.000 | 0.000 |
| 48 | *Sphyrna_lewini* | *Sphyrna_lewini* | Match | 0.785 | 0.000 | 0.000 | 0.009 | 0.000 | 0.005 | 0.000 | 0.004 | 0.000 | 0.001 | 0.000 | 0.000 | 0.001 | 0.000 | 0.000 | 0.021 | 0.009 | 0.156 | 0.000 | 0.000 | 0.000 | 0.000 | 0.000 | 0.000 | 0.000 | 0.785 | 0.005 | 0.003 | 0.000 |
| 49 | *Sphyrna_lewini* | *Glaucostegus_typus* | Mismatch | 0.914 | 0.000 | 0.000 | 0.000 | 0.000 | 0.000 | 0.000 | 0.000 | 0.000 | 0.914 | 0.000 | 0.000 | 0.001 | 0.000 | 0.000 | 0.000 | 0.001 | 0.000 | 0.000 | 0.000 | 0.000 | 0.000 | 0.000 | 0.000 | 0.001 | 0.083 | 0.000 | 0.000 | 0.000 |
| 50 | *Sphyrna_lewini* | *Sphyrna_lewini* | Match | 0.987 | 0.000 | 0.000 | 0.000 | 0.000 | 0.008 | 0.000 | 0.000 | 0.000 | 0.001 | 0.000 | 0.000 | 0.000 | 0.000 | 0.000 | 0.000 | 0.000 | 0.002 | 0.001 | 0.000 | 0.000 | 0.000 | 0.000 | 0.000 | 0.000 | 0.987 | 0.000 | 0.000 | 0.000 |
| 51 | *Alopias_superciliosus* | *Alopias_superciliosus* | Match | 0.608 | 0.000 | 0.608 | 0.000 | 0.000 | 0.000 | 0.000 | 0.000 | 0.000 | 0.000 | 0.000 | 0.000 | 0.000 | 0.000 | 0.000 | 0.000 | 0.000 | 0.000 | 0.000 | 0.000 | 0.000 | 0.000 | 0.000 | 0.392 | 0.000 | 0.000 | 0.000 | 0.000 | 0.000 |
| 52 | *Carcharhinus_longimanus* | *Glaucostegus_typus* | Mismatch | 1.000 | 0.000 | 0.000 | 0.000 | 0.000 | 0.000 | 0.000 | 0.000 | 0.000 | 1.000 | 0.000 | 0.000 | 0.000 | 0.000 | 0.000 | 0.000 | 0.000 | 0.000 | 0.000 | 0.000 | 0.000 | 0.000 | 0.000 | 0.000 | 0.000 | 0.000 | 0.000 | 0.000 | 0.000 |
| 53 | *Carcharhinus_longimanus* | *Glaucostegus_typus* | Mismatch | 1.000 | 0.000 | 0.000 | 0.000 | 0.000 | 0.000 | 0.000 | 0.000 | 0.000 | 1.000 | 0.000 | 0.000 | 0.000 | 0.000 | 0.000 | 0.000 | 0.000 | 0.000 | 0.000 | 0.000 | 0.000 | 0.000 | 0.000 | 0.000 | 0.000 | 0.000 | 0.000 | 0.000 | 0.000 |
| 54 | *Carcharhinus_longimanus* | *Glaucostegus_typus* | Mismatch | 1.000 | 0.000 | 0.000 | 0.000 | 0.000 | 0.000 | 0.000 | 0.000 | 0.000 | 1.000 | 0.000 | 0.000 | 0.000 | 0.000 | 0.000 | 0.000 | 0.000 | 0.000 | 0.000 | 0.000 | 0.000 | 0.000 | 0.000 | 0.000 | 0.000 | 0.000 | 0.000 | 0.000 | 0.000 |
| 55 | *Isurus_oxyrinchus* | *Isurus_oxyrinchus* | Match | 1.000 | 0.000 | 0.000 | 0.000 | 0.000 | 0.000 | 0.000 | 0.000 | 0.000 | 0.000 | 0.000 | 0.000 | 1.000 | 0.000 | 0.000 | 0.000 | 0.000 | 0.000 | 0.000 | 0.000 | 0.000 | 0.000 | 0.000 | 0.000 | 0.000 | 0.000 | 0.000 | 0.000 | 0.000 |
| 56 | *Sphyrna_mokarran* | *Sphyrna_mokarran* | Match | 0.973 | 0.000 | 0.000 | 0.000 | 0.000 | 0.000 | 0.015 | 0.000 | 0.000 | 0.001 | 0.000 | 0.000 | 0.000 | 0.001 | 0.003 | 0.007 | 0.000 | 0.000 | 0.000 | 0.000 | 0.000 | 0.000 | 0.000 | 0.000 | 0.000 | 0.000 | 0.973 | 0.000 | 0.000 |
| 57 | *Stegostoma_fasciatum* | *Stegostoma_fasciatum* | Match | 0.523 | 0.000 | 0.000 | 0.000 | 0.000 | 0.000 | 0.000 | 0.477 | 0.000 | 0.000 | 0.000 | 0.000 | 0.000 | 0.000 | 0.000 | 0.000 | 0.000 | 0.000 | 0.000 | 0.000 | 0.000 | 0.000 | 0.000 | 0.000 | 0.000 | 0.000 | 0.000 | 0.523 | 0.000 |
| 58 | *Alopias_pelagicus* | *Alopias_pelagicus* | Match | 1.000 | 1.000 | 0.000 | 0.000 | 0.000 | 0.000 | 0.000 | 0.000 | 0.000 | 0.000 | 0.000 | 0.000 | 0.000 | 0.000 | 0.000 | 0.000 | 0.000 | 0.000 | 0.000 | 0.000 | 0.000 | 0.000 | 0.000 | 0.000 | 0.000 | 0.000 | 0.000 | 0.000 | 0.000 |
| 59 | *Anoxypristis_cuspidata* | *Anoxypristis_cuspidata* | Match | 1.000 | 0.000 | 0.000 | 1.000 | 0.000 | 0.000 | 0.000 | 0.000 | 0.000 | 0.000 | 0.000 | 0.000 | 0.000 | 0.000 | 0.000 | 0.000 | 0.000 | 0.000 | 0.000 | 0.000 | 0.000 | 0.000 | 0.000 | 0.000 | 0.000 | 0.000 | 0.000 | 0.000 | 0.000 |
| 60 | *Lamna_nasus* | *Lamna_nasus* | Match | 1.000 | 0.000 | 0.000 | 0.000 | 0.000 | 0.000 | 0.000 | 0.000 | 0.000 | 0.000 | 0.000 | 0.000 | 0.000 | 0.000 | 1.000 | 0.000 | 0.000 | 0.000 | 0.000 | 0.000 | 0.000 | 0.000 | 0.000 | 0.000 | 0.000 | 0.000 | 0.000 | 0.000 | 0.000 |
| 61 | *Lamna_nasus* | *Lamna_nasus* | Match | 1.000 | 0.000 | 0.000 | 0.000 | 0.000 | 0.000 | 0.000 | 0.000 | 0.000 | 0.000 | 0.000 | 0.000 | 0.000 | 0.000 | 1.000 | 0.000 | 0.000 | 0.000 | 0.000 | 0.000 | 0.000 | 0.000 | 0.000 | 0.000 | 0.000 | 0.000 | 0.000 | 0.000 | 0.000 |
| 62 | *Mobula_birostris* | *Mobula_birostris* | Match | 1.000 | 0.000 | 0.000 | 0.000 | 0.000 | 0.000 | 0.000 | 0.000 | 0.000 | 0.000 | 0.000 | 0.000 | 0.000 | 0.000 | 0.000 | 1.000 | 0.000 | 0.000 | 0.000 | 0.000 | 0.000 | 0.000 | 0.000 | 0.000 | 0.000 | 0.000 | 0.000 | 0.000 | 0.000 |
| 63 | *Mobula_mobular* | *Mobula_mobular* | Match | 1.000 | 0.000 | 0.000 | 0.000 | 0.000 | 0.000 | 0.000 | 0.000 | 0.000 | 0.000 | 0.000 | 0.000 | 0.000 | 0.000 | 0.000 | 0.000 | 1.000 | 0.000 | 0.000 | 0.000 | 0.000 | 0.000 | 0.000 | 0.000 | 0.000 | 0.000 | 0.000 | 0.000 | 0.000 |
| 64 | *Mobula_tarapacana* | *Mobula_tarapacana* | Match | 1.000 | 0.000 | 0.000 | 0.000 | 0.000 | 0.000 | 0.000 | 0.000 | 0.000 | 0.000 | 0.000 | 0.000 | 0.000 | 0.000 | 0.000 | 0.000 | 0.000 | 1.000 | 0.000 | 0.000 | 0.000 | 0.000 | 0.000 | 0.000 | 0.000 | 0.000 | 0.000 | 0.000 | 0.000 |
| 65 | *Pristis_pristis* | *Pristis_pristis* | Match | 1.000 | 0.000 | 0.000 | 0.000 | 0.000 | 0.000 | 0.000 | 0.000 | 0.000 | 0.000 | 0.000 | 0.000 | 0.000 | 0.000 | 0.000 | 0.000 | 0.000 | 0.000 | 0.000 | 0.000 | 1.000 | 0.000 | 0.000 | 0.000 | 0.000 | 0.000 | 0.000 | 0.000 | 0.000 |
| 66 | *Carcharhinus_falciformis* | *Carcharhinus_falciformis* | Match | 1.000 | 0.000 | 0.000 | 0.000 | 0.000 | 1.000 | 0.000 | 0.000 | 0.000 | 0.000 | 0.000 | 0.000 | 0.000 | 0.000 | 0.000 | 0.000 | 0.000 | 0.000 | 0.000 | 0.000 | 0.000 | 0.000 | 0.000 | 0.000 | 0.000 | 0.000 | 0.000 | 0.000 | 0.000 |
| 67 | *Carcharhinus_falciformis* | *Carcharhinus_falciformis* | Match | 1.000 | 0.000 | 0.000 | 0.000 | 0.000 | 1.000 | 0.000 | 0.000 | 0.000 | 0.000 | 0.000 | 0.000 | 0.000 | 0.000 | 0.000 | 0.000 | 0.000 | 0.000 | 0.000 | 0.000 | 0.000 | 0.000 | 0.000 | 0.000 | 0.000 | 0.000 | 0.000 | 0.000 | 0.000 |
| 68 | *Carcharhinus_falciformis* | *Carcharhinus_falciformis* | Match | 1.000 | 0.000 | 0.000 | 0.000 | 0.000 | 1.000 | 0.000 | 0.000 | 0.000 | 0.000 | 0.000 | 0.000 | 0.000 | 0.000 | 0.000 | 0.000 | 0.000 | 0.000 | 0.000 | 0.000 | 0.000 | 0.000 | 0.000 | 0.000 | 0.000 | 0.000 | 0.000 | 0.000 | 0.000 |

**Table S.6**. Initial value of hyper-parameters in searching for the best deep learning model using grid search method

| Parameters | Definition | Value |
| --- | --- | --- |
| activation | The activation function of learning model | "Rectifier", "Maxout", "Tanh", "RectifierWithDropout", "MaxoutWithDropout" and "TanhWithDropout" |
| hidden | Number of learning layers | [100, 100, 100], [200, 200, 200] and [500, 500, 500] |
| epochs | Number of times to iterate (stream) the dataset | 50, 100, 200, 300 and 500 |
| rho | The adaptive learning rate time decay factor | 0.9, 0.95, 0.99 and 0.999 |
| epsilon | The adaptive learning rate time smoothing factor to avoid dividing by zero | 1e-10, 1e-8, 1e-6 and 1e-4 |
| input_dropout_ratio | The input layer dropout ratio to improve generalisation. Suggested values are 0.1 or 0.2 | 0, 0.1 and 0.2 |
| l1 | The L1 regularization to add stability and improve generalisation | 0, 0.00001 and 0.0001 |
| l2 | The L2 regularization to add stability and improve generalisation | 0, 0.00001 and 0.0001 |
| max_w2 | The constraint for the squared sum of the incoming weights per unit | 10, 100, 1000 and 3.4028235e+38 |

**Table S.7**. Stopping criteria in searching the best deep learning model

| Criteria | Definition | Value |
|---|---|---|
| strategy | strategy to perform a random search of all the combinations of your hyperparameters | RandomDiscrete |
| max_models | The maximum number of generated models | 100,000 |
| max_runtime_secs | The maximum run time in second | 43,200 seconds (12 hours) |
| stopping_tolerance | Stop if MSE hasn't improved by the value | 0.001 |
| stopping_rounds | Number of models to compare MSE improvement | 20 |
| seed | Seed number to control randomness | 1234 |
